# Engineering site-specific nucleic acid-protein conjugates by utilizing a natural RNAylation reaction

**DOI:** 10.1101/2025.07.11.664405

**Authors:** Nurseda Yilmaz Demirel, Ilayda Mumcuoglu, Gabriele Malengo, Bernd Schmeck, Anna Lena Jung, Katharina Höfer

**Affiliations:** Max Planck Institute for Terrestrial Microbiology, Marburg, Germany; Center for Synthetic Microbiology (SYNMIKRO), Philipps Universität Marburg, Marburg, Germany; Institute for Lung Research, Philipps Universität Marburg, Marburg, Germany; German Center for Lung Research (DZL), Giessen, Germany; Institute for Lung Health (ILH), Justus-Liebig University, Giessen, Germany; Department of Medicine, Pulmonary and Critical Care Medicine, University Hospital Giessen and Marburg, Philipps Universität Marburg, Marburg, Germany; Core Facility Flow Cytometry - Bacterial Vesicles, Philipps Universität Marburg, Marburg, Germany; Department of Pharmacy, Institute of Pharmaceutical Biology and Biotechnology, Philipps Universität Marburg, Marburg, Germany

## Abstract

Nucleic acid-protein conjugates are valuable for synthetic biology, therapeutics, and nanotechnology, but current methods often lack site specificity and rely on non-natural linkages. RNAylation, a one-step enzymatic reaction catalyzed by the bacteriophage T4 enzyme ModB where first discovered *in vivo* during phage infection, enables site-specific conjugation of nucleic acids to proteins via a natural N-glycosidic bond. Here, we establish RNAylation as a novel and robust *in vitro* platform for generating nucleic acid-protein conjugates, overcoming key limitations of existing strategies. We define design principles for this approach, demonstrate enhanced nucleic acid stability in human cell lysates, and develop an efficient purification workflow. Furthermore, we achieve successful delivery of purified conjugates into human cells, highlighting the potential for functional *in vivo* applications. Our work expands RNAylation from a phage-specific phenomenon to a versatile, biologically relevant strategy with broad biotechnological potential.

## Introduction

Nucleic acids and proteins are fundamental functional building blocks of life, each with distinct biological properties. Nucleic acids, such as RNA and DNA, possess high programmability and structural predictability due to specific base-pairing interactions. Additionally, nucleic acids serve as carriers of genetic information, properties that make them valuable for genetic engineering, therapeutics, diagnostics, and nanotechnology^1-3^. In contrast, proteins exhibit remarkable structural and functional diversity, making them indispensable for the development of molecular tools and systems. Nature has demonstrated that interactions between nucleic acids and proteins can give rise to highly important cellular machineries, such as the ribosome^4^. This complex, which is central to translation, exemplifies how the synergy between nucleic acids and proteins can drive critical biological functions. Notably, these natural assemblies are typically stabilized by transient, non-covalent interactions, allowing for dynamic regulation and adaptability within the cell^5^.

Historically, nucleic acids and proteins were studied and applied independently. However, recognizing the significance of naturally occurring nucleic acid-protein assemblies, researchers have sought to harness and enhance their synergistic properties by designing and constructing nucleic acid-protein conjugates *in vitro*^6, 7^. Applications of these conjugates now span targeted drug delivery, genome editing, biosensing, nanostructure assembly, and the precise manipulation of cellular processes. RNA–protein conjugates are particularly valuable, combining RNA’s programmability and information-carrying capacity with protein functionality, resulting in biomolecules with enhanced characteristics. Such conjugates can improve RNA stability against nucleases and extend RNA’s functional lifetime in biological environments^8^. They also enable the design of programmable scaffolds, sensors, and delivery systems, where nucleic acids provide targeting or regulatory functions and proteins contribute catalytic or binding activities^9-11^.

To mimic nature and to generate nucleic-acid conjugates *in vitro*, several biochemical and chemical strategies were developed in the past. Chemical conjugation strategies, such as lysine or cysteine modification, often lack site selectivity due to the abundance and accessibility of reactive sites in a single protein^12, 13^, while approaches using non-canonical amino acids or click chemistry are labor-intensive and costly^14, 15^. Moreover, these methods typically form non-natural linkages, raising concerns about the toxicity of the produced conjugates in biological systems. Biochemical methods, including self-ligating protein tags like SNAP-tags, offer site specificity but require specialized nucleic acid substrates, increasing system complexity^16, 17^. Moreover, the SNAP-tag forms a highly stable thioether bond with its substrate, a linkage for which no natural enzymes are known to mediate cleavage^18^. As a result, these conjugates are exceptionally resistant to degradation in biological systems, potentially increasing the risk of immunogenic responses *in vivo*. Despite significant progress in the generation of nucleic acid-protein conjugates, existing approaches continue to face persistent challenges, including limited site specificity, labor-intensive protocols, and the formation of non-natural linkages that may compromise biocompatibility. Thus, there is a need for efficient, site-specific, and biocompatible conjugation technologies that remain unmet.

Here, we present a versatile conjugation strategy that enables site-specific linkage of virtually any nucleic acid to any protein via an RNAylation-based mechanism. RNAylation is a one-step enzymatic process originally discovered during bacteriophage T4 infection of *Escherichia coli*, in which the ADP-ribosyltransferase ModB transfers an ADP-ribose-RNA (ADPr-RNA) moiety from nicotinamide adenine dinucleotide (NAD^+^)-capped RNA (NAD-RNA) to specific arginine residues on target proteins^19^. This reaction forms a covalent N-glycosidic bond with high site selectivity. To date, the primary targets of ModB-mediated RNAylation are ribosomal proteins S1 (rS1) and L2 (rL2)^19^, both RNA-binding proteins with accessible arginine residues in their nucleic acid-binding domains^20-23^. To date, NAD-capped RNAs have been identified as the sole substrates for ModB-mediated RNAylation^19^. First discovered in *E. coli*^24, 25^, NAD-RNAs are now recognized across diverse organisms^26-36^, underscoring their evolutionary conservation and biological relevance.

Inspired by the ability of ModB to generate RNA-protein conjugates based on a natural bond with site-selectivity, we introduce RNAylation as a novel tool to generate RNA-protein conjugates and expand its utility in the field of synthetic biology. In this study, we broaden the substrate scope of ModB by introducing non-canonical substrates: NGD-, NCD-, and NUD-RNAs, collectively termed NXD-RNAs. Moreover, for the first time, we demonstrate DNAylation, showing that NXD-DNAs can also be covalently linked to proteins. This advancement enables the covalent attachment of diverse NXD-capped nucleic acids to proteins, greatly enhancing the flexibility and utility of the platform. Importantly, the bond formed by ModB between nucleic acids and proteins is a naturally occurring N-glycosidic linkage, which can be hydrolyzed by eukaryotic enzymes such as ADP-ribosylhydrolase 1 (ARH1)^19^. The ability to generate conjugates with a cleavable, biologically recognized bond not only facilitates controlled release and recycling but also aligns with natural cellular processes, enhancing the biocompatibility of these constructs in diverse applications. To further expand the versatility of our platform, we engineered an RNAylation tag that enables consistent and site-specific modification in a standardized way. The designed RNAylation tag is a fusion of the nucleic acid binding domain II of rS1 protein^37^ and SpyTag, establishing a modular system based on third-generation SpyTag/SpyCatcher technology^38^ that enables the customized assembly of diverse nucleic acid–protein conjugates. Assessing the stability and deliverability of these conjugates is crucial; our results show that covalent attachment of RNA to a protein enhances RNA stability in human cell lysates, and purified DNA–protein conjugates are successfully delivered into human cells. Thereby, the application of RNAylation can provide a versatile platform for creating dual biomolecules with synergistic properties, paving the way for advances in programmable molecular scaffolds, enhanced molecular stability, targeted delivery systems, and nucleic acid therapeutics.

## Results

### Versatile RNA–protein conjugate generation mediated by ModB

ModB-mediated RNAylation can be utilized as a synthetic biology tool to generate RNA-protein conjugates. To unlock the full potential of RNAylation to generate novel nucleic acid-protein conjugates with unique properties, we systematically investigated the substrate scope of ModB. Historically, NAD^39^ and NAD-RNAs^19^ are the only defined biological substrates for ModB. Mechanistic investigations of ARTs indicate that the nicotinamide mononucleotide (NMN) moiety of NAD is essential for catalysis, serving as the primary determinant for substrate recognition and enzymatic activity^40^. This observation is further supported by studies employing modified NAD derivatives, such as clickable NAD (2-Ethynyl-Adenosine-NAD, 6-alkyne-NAD, 8-alkyne-NAD)^41, 42^, fluorescein-NAD^43^, biotin-NAD^44^, and Etheno-NAD^45^, in studying protein ADP-ribosylation by ARTs^46^. Remarkably, the mentioned modifications are consistently localized to the adenosine residue, while the nicotinamide mononucleotide (NMN) moiety remains unaltered, highlighting its conserved role for the reactions.

Consistent with these biochemical observations, we employed AlphaFold3 prediction^47^ of ModB (Figure 1 and Supplementary Figure 1) bound to NAD, revealing that the NMN moiety is positioned between the glutamic acid at position 173 (E173) and arginine at position 73 (R73) with high confidence of prediction. It has been shown that E173 is essential for enzymatic activity, while R73 plays a crucial role in NAD binding to facilitate ADP-ribosylation^48^. On the other hand, adenine is located further from the catalytic site, suggesting that it may have less influence on enzymatic function.

**Figure 1:**
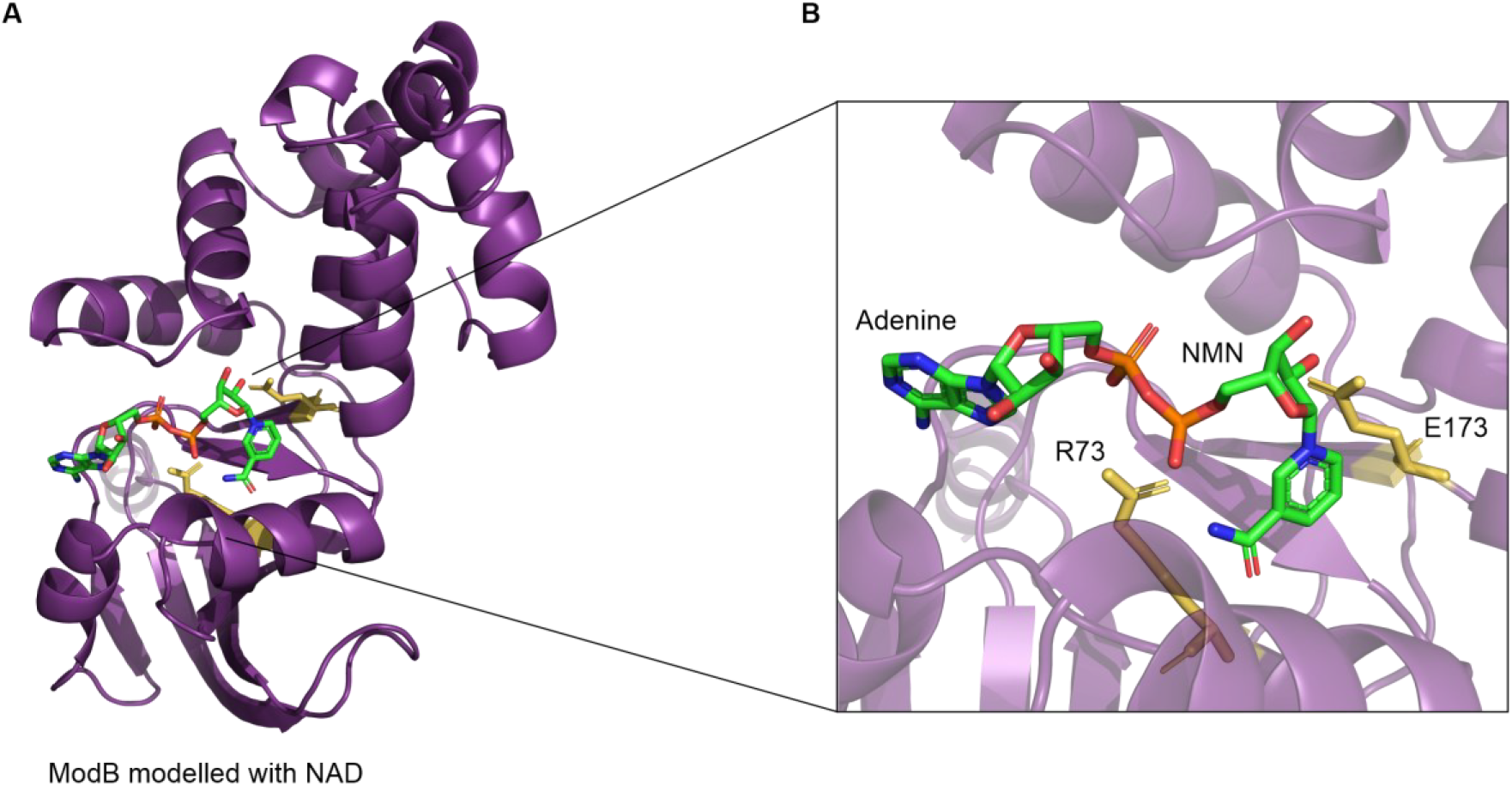
AlphaFold3 structural prediction of ModB bound with NAD substrate. A) A structural model of ModB enzyme bound to its natural substrate NAD is predicted by AlphaFold3^47^ with the predicted Template Modeling score (pTM) of 0.89, showing high confidence. B) The zoomed-in catalytic center demonstrates the substrate binding and highlights key residues for enzymatic activity. NMN is located between the residues R73 and E173, which are crucial for substrate binding and catalytic activity, respectively^48^.

Given that RNAylation follows an ADP-ribosylation-like mechanism, the NAD moiety of NAD-RNA is likely to be positioned similarly, with the RNA side chain extended from the adenine. Based on these observations, we hypothesize that the adenine moiety of NAD-RNA can be replaced by other bases such as guanine, cytosine, and uracil without hampering the enzymatic activity of ModB. To test our hypothesis, we first synthesized NGD-, NDC-, and NUD-capped RNAs based on the published phosphorimidazolide chemistry-based approach^49^ (Supplementary Figure 2). Collectively, all capped RNAs (NAD-, NGD-, NCD-, and NUD-RNAs) are named as NXD-RNAs. The capping efficiencies for NXD-RNAs are in agreement with the published data for NAD-RNA, which range between 23% and 30% (Supplementary Figure 2). To test our hypothesis that NXD-RNAs can be attached to a protein, the described target protein rS1 was subjected to reactions mediated by ModB in the presence of NXD-RNAs. All tested NXD-RNAs were successfully attached to rS1, as demonstrated by gel electrophoresis using 3′ Cy5-labeled NXD-RNAs (Figure 2A, Supplementary Figure 3). NAD-RNA served as a positive control, while 5′-phosphorylated-X-RNAs (P-X-RNAs) (lacking a cap) served as negative controls. No RNAylation was observed in the absence of ModB or with P-X-RNAs, confirming that conjugation is both enzyme-and cap-dependent. Notably, RNAylation yields were enhanced with NGD-RNA and NUD-RNA (2-and 2.6-fold, respectively) relative to NAD-RNA, while NCD-RNA yielded a fourfold increase compared to NAD-RNA (Figure 2B). Our results demonstrate that ModB can RNAylate rS1 regardless of the identity of the nucleobase in the cap structure, establishing NGD-RNA, NCD-RNA, and NUD-RNA as novel substrates for ModB.

**Figure 2:**
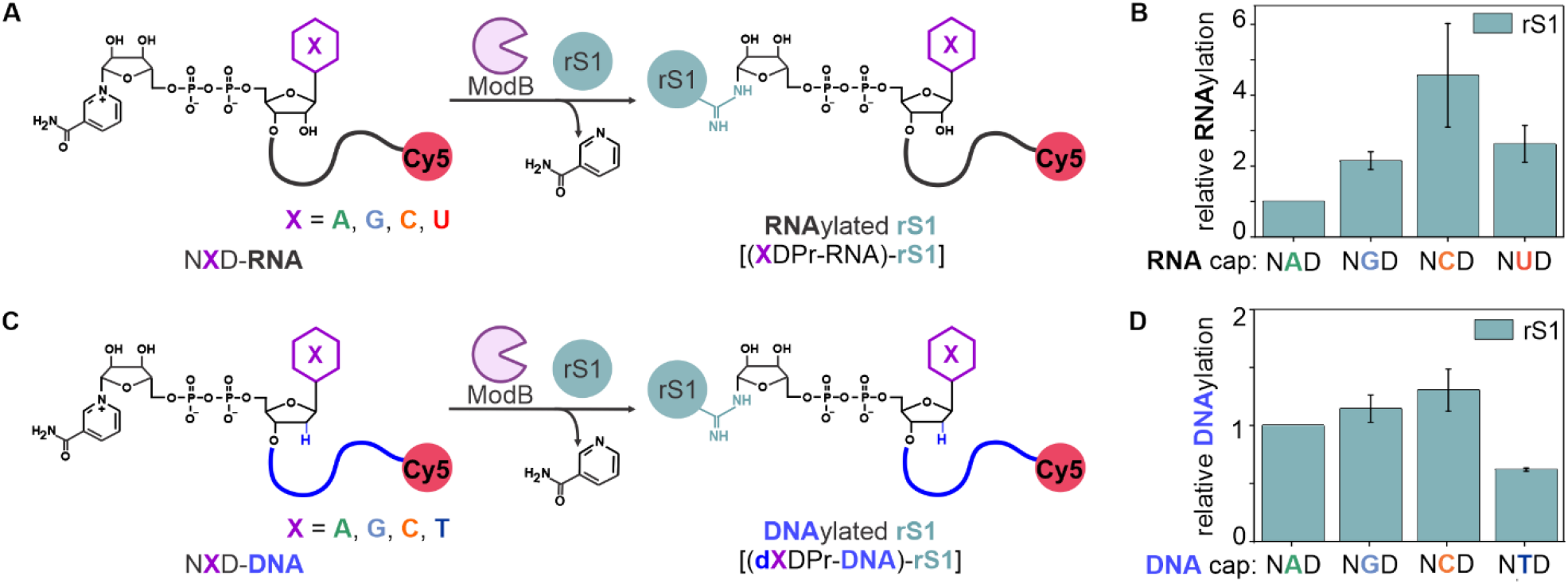
The covalent attachment of NXD-RNAs and NXD-DNAs to rS1 by ModB. A) The proposed RNAylation-reaction mechanism of rS1 in the presence of NXD-RNAs is depicted. B) Relative RNAylation efficiencies of rS1 with NXD-RNAs have been shown. Relative RNAylation yields were calculated using NAD-RNA as the standard. Reactions have been performed in the presence of 3’ Cy5 labelled NXD-RNA 10mers. Error bars represent the mean ± s.d. of n = 3 independent experiments. C) The proposed DNAylation-reaction mechanism of rS1 in the presence of NXD-DNAs is demonstrated. The same reaction mechanism as RNAylation is hypothesized for the DNAylation reaction. D) Relative rS1 DNAylation efficiencies in the presence of NXD-DNAs are presented. Relative DNAylation yields were calculated using NAD-DNA as the standard. Reactions have been performed in the presence of 3’ Cy5 labelled NXD-DNA 40mers. Error bars represent the mean ± s.d. of n = 3 independent experiments.

### ModB-driven DNAylation facilitates DNA-protein conjugate generation

The successful use of NXD-RNAs indicates that the nicotinamide moiety is essential for substrate recognition, whereas the adenine base can be substituted without loss of activity. Building on this, we hypothesize that NXD-capped DNAs (NXD-DNAs) could also serve as substrates for ModB. After synthesizing NXD-DNAs *in vitro* (Supplementary Figure 4), we demonstrated their covalent attachment to rS1 (Supplementary Figure 5) via ModB, establishing the first example of ModB-mediated DNA-protein conjugate generation, which we term DNAylation. This finding expands the substrate scope of ModB and introduces a new strategy for site-specific DNA-protein conjugate formation, thereby broadening the utility of our RNAylation platform to include diverse nucleic acids. Interestingly, we observe a different trend for DNAylation than RNAylation. While NAD-, NGD-, and NCD-DNAs yielded similar DNAylation efficiency, NTD-DNA led to a 40% efficiency decrease (Figure 2D). Collectively, these findings establish that ModB can covalently link a wide range of capped nucleic acids—both RNAs and DNAs—to proteins in a single-step enzymatic reaction. These new design principles position ModB as a versatile synthetic biology tool for the site-specific conjugation of diverse nucleic acid substrates to proteins.

### Modular system establishment for flexible nucleic acid–protein conjugate generation

To facilitate the flexible generation of nucleic acid–protein conjugates for diverse applications, we established a modular system that enables the use of any protein of interest (POI) as a conjugation partner. While expanding the substrate scope of ModB allows the use of any RNA or DNA sequence, the natural system is limited to just two identified target proteins. To overcome this limitation, we engineered a strategy based on the small rS1 domain II (DII), a 10.7 kDa fragment previously shown to be efficiently RNAylated *in vitro*^19^, as a universal tag that can be fused to any POI. We demonstrated that NXD-RNAs can be covalently attached to rS1 DII with yields comparable to those observed for full-length rS1 (Supplementary Figure 6).

Moreover, to prove our hypothesis that any POI can be RNAylated by the fusion of rS1 DII as a tag, we have generated the fusion protein of GFP to rS1 DII (GFP-DII). We have subjected GFP and GFP-DII to ModB-mediated RNAylation (Supplementary Figure 7). We demonstrated that the RNAylation of GFP-DII by visualizing a signal in the Cy5 channel only in the presence of NAD-RNA and ModB (Supplementary Figure 7B). RNAylated GFP-DII is shifted upwards on the gel compared to unmodified GFP-DII since the covalent attachment of RNA increases the molecular weight of the protein. On the other hand, we were able to show that NAD-RNA cannot be attached to untagged GFP, confirming the tag’s specificity and utility. Therefore, rS1 DII can serve as an RNAylation tag to covalently link an RNA with a POI.

To further enhance modularity and to ensure RNAylation/DNAylation of specific arginine residues, we applied SpyCatcher-SpyTag technology. The SpyCatcher-SpyTag system is designed based on a modified surface protein from *Streptococcus pyogenes* (SpyCatcher) and a 13 amino acid peptide tag (SpyTag) that can be recognized by SpyCatcher^50, 51^. When SpyCatcher and SpyTag interact, an irreversible covalent bond is formed between them. To employ this technology, we generated a SpyTag–rS1 DII fusion (SpyTag-DII) as an RNAylation tag. Initial experiments revealed that RNAylated SpyTag-DII cannot form a bond with SpyCatcher-GFP as efficiently as unmodified SpyTag-DII (Figure 3A). Analysis using liquid chromatography with tandem mass spectrometry (LC–MS/MS) has shown that an arginine residue of the SpyTag part is RNAylated when SpyTag-DII is subjected to an RNAylation reaction (Supplementary Figure 8). To address this, we created a mutant RNAylation tag (SpyTag R3K-DII), substituting the arginine with lysine to maintain charge while preventing unwanted RNAylation at that site. The utilization of a mutated RNAylation tag restored efficient conjugation between RNAylated tag and SpyCatcher-GFP, establishing a robust and versatile modular system (Figure 3B). Therefore, in this study, we have engineered SpyTag R3K-DII as an RNAylation tag that can be modified at specific residues. Collectively, these results demonstrate that our approach enables the flexible and site-specific conjugation of any protein of interest to any nucleic acid sequence, significantly broadening the utility of ModB-mediated conjugation for synthetic biology and biotechnological applications.

**Figure 3:**
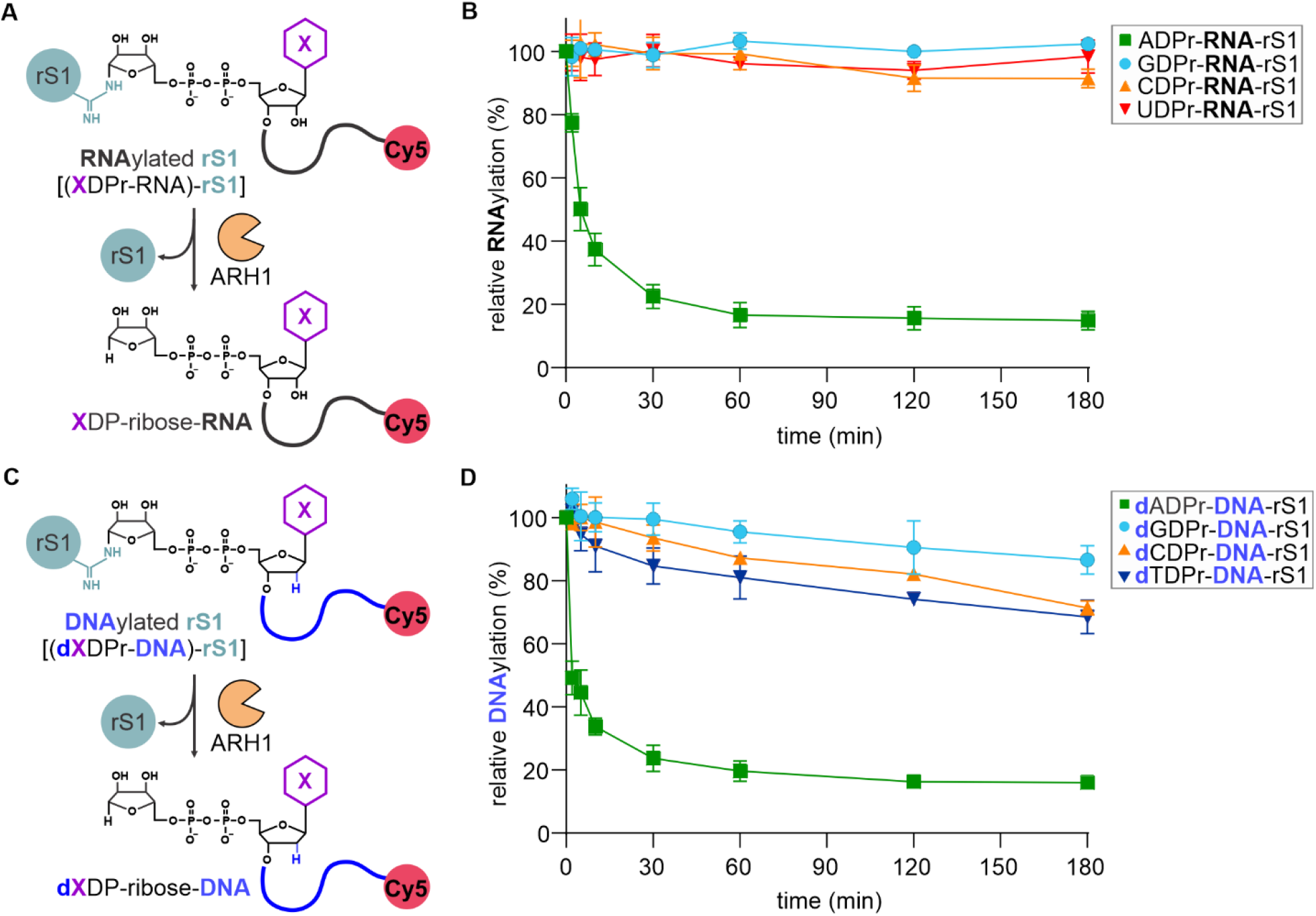
Establishment of modular system based on SpyCatcher-SpyTag technology. A) A schematic representation of the modular system is demonstrated on the left side. The reactions of SpyCatcher-GFP with RNAylation tag (SpyTag-DII) or RNAylated RNAylation tag are analyzed via 12% SDS-PAGE. For RNAylation reaction, 3’Cy5 labelled NAD-RNA 10mer is used. (n = 3 independent experiments) B) Schematic representation of the modular system with newly engineered RNAylation tag* (SpyTag R3K-DII) is demonstrated on the left side. The reactions of SpyCatcher-GFP with RNAylation tag* (SpyTag R3K-DII) or RNAylated RNAylation tag* are analyzed via 12% SDS-PAGE. For RNAylation reaction, 3’Cy5 labelled NAD-RNA 10mer is used. (n = 3 independent experiments)

### Evaluation of the stability of nucleic-acid protein conjugates against ARH1

The eukaryotic enzyme ADP-ribosylhydrolase 1 (ARH1) specifically cleaves the N-glycosidic linkage of ADPr-RNA-arginine residues in RNAylated proteins^19^. Mechanistic studies have shown that ARH1 recognizes the adenosine moiety of the attached ADP-ribose, forming direct interactions with key protein residues (Ser124, Gly127, and Ser270), which are essential for substrate positioning within the catalytic center^52^. Based on these insights, we hypothesize that substituting the adenine base in the NAD cap structure with alternative nucleobases (guanine, cytosine, or uracil) might disrupt ARH1 recognition and thereby enhance the stability of the nucleic acid-protein conjugates. To test this hypothesis, we conducted ARH1 cleavage assays using rS1 conjugated to various NXD-capped RNAs (where X = G, C, or U) and compared them to the naturally occurring ADPr-RNA-rS1 conjugate (Supplementary Figure 9). Our results revealed that ARH1 has hydrolyzed 80% of the N-glycosidic bond of ADPr-RNA-rS1 within 30 minutes (Figure 4A) *in vitro*. On the other hand, GDPr-, CDPr-, UDPr-RNA-rS1 conjugates remained mostly intact even after 3 hours incubation with at least 90% of RNAylated rS1 still detected (Figure 4A). We extended this analysis to DNAylated conjugates, reasoning that ARH1 would show similar substrate preferences for dXDPr-DNA-rS1 conjugates. Consistent with our findings for RNAylated conjugates, 80% of dADPr-DNA was cleaved by ARH1 within 30 minutes, whereas dGDPr-, dCDPr, and dTDPr-DNA-rS1 conjugates exhibited pronounced stability, with only up to 20% cleavage observed for dTDPr-DNA-rS1 after 3 hours (Supplementary Figure 10, Figure 4B).

Collectively, these results indicate that ARH1 exhibits a strong preference for ADPr-linkages, and its enzymatic activity is substantially diminished when adenine is replaced by other nucleobases *in vitro*. Thus, the stability of RNAylated and DNAylated proteins can be significantly enhanced against ARH1 by employing non-canonical capped nucleic acids as substrates for ModB. This strategy enables the rational design of conjugates with tailored resistance to enzymatic cleavage, broadening their potential for both *in vitro* and *in vivo* applications where conjugate stability is paramount.

**Figure 4:**
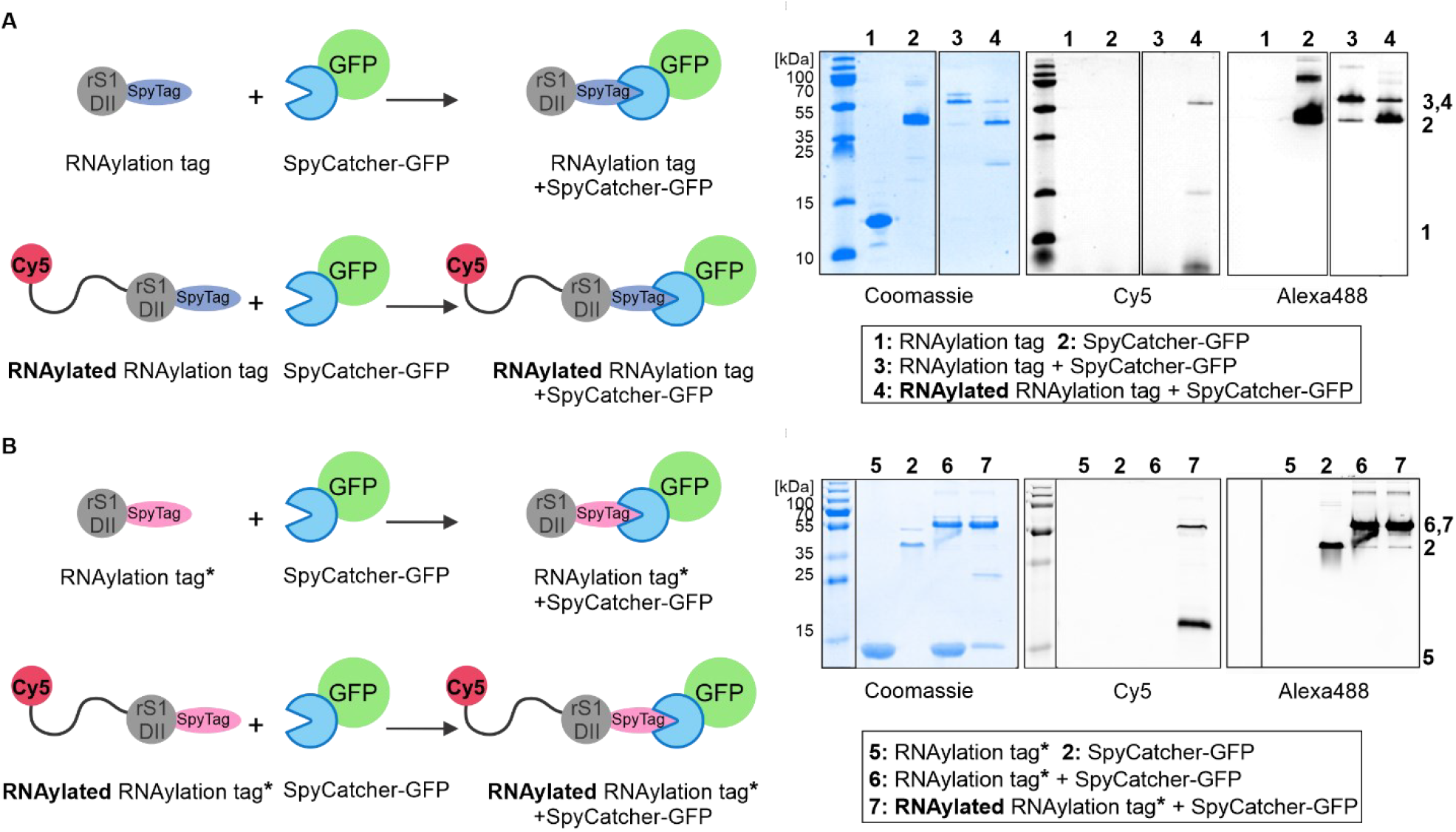
NXD-RNAylated rS1 and NXD-DNAylated rS1 stability against ARH1. A) The described reaction mechanism of ARH1 on RNAylated protein has been shown. We hypothesize that ARH1 activity will be hampered when adenine is exchanged with other bases. B) ARH1 kinetics graph is shown for NXD-RNAylated rS1, where RNAylation is performed in the presence of 3’ Cy5 labelled NXD-RNA 10mers. Error bars represent the mean ± s.d. of n = 3 independent experiments. C) The same reaction mechanism as the cleavage of dXDPr-DNA from rS1 is predicted for NXD-DNAylated rS1. D) ARH1 kinetics graph of NXD-DNAylated rS1, where DNAylation is performed in the presence of 3’ Cy5 labelled NXD-DNA 40mers, is demonstrated. Error bars represent the mean ± s.d. of n = 3 independent experiments.

### Assessment of the stability of RNAylated proteins under physiological conditions

A key factor for the practical utility of RNAylated proteins is their stability, particularly in environments where nucleic acids are prone to degradation. To assess whether covalent attachment of RNA to a protein enhances RNA stability against exonucleases, we evaluated the resistance of NXD-RNAylated rS1 to XRN1^53^, the major cytoplasmic 5′-3′ exoribonuclease involved in mRNA decay in eukaryotic cells. Using XRN1 kinetics assays, we compared the degradation of NXD-RNAylated rS1 with that of 5′-phosphorylated adenosine RNA (5′-P-A-RNA), a canonical XRN1 substrate, as a positive control (Supplementary Figure 11). After 3 hours of XRN1 treatment, only 15% of the initial 5′-P-A-RNA remained, confirming its rapid degradation. In contrast, at least 80% of the NXD-RNAylated rS1 conjugates persisted after the same incubation time, demonstrating that covalent linkage to a protein at the 5′ end effectively shields RNA from XRN1-mediated cleavage. This result highlights the protective effect of protein conjugation against exonuclease activity *in vitro*.

Building on these findings, we examined the stability of NXD-RNAylated conjugates in more complex biological environments to mimic *in vivo* conditions. To test the stability of RNA-protein conjugates, we utilized a target protein rL2 fused with GFP (GFP-rL2). In human cell lysate (prepared from human embryonic kidney (HEK293T) cells), all tested NXD-RNAylated GFP-rL2 conjugates remained stable over a 24-hour time course, with 50% of the initial conjugate still detectable after 24 hours (Figure 5A, Supplementary Figure 12A-E). In contrast, 5′-P-A-RNA was rapidly degraded in the same lysate, with only 3% remaining after one hour. These results confirm that covalent attachment of RNA to a protein significantly enhances RNA stability under cellular conditions.

**Figure 5:**
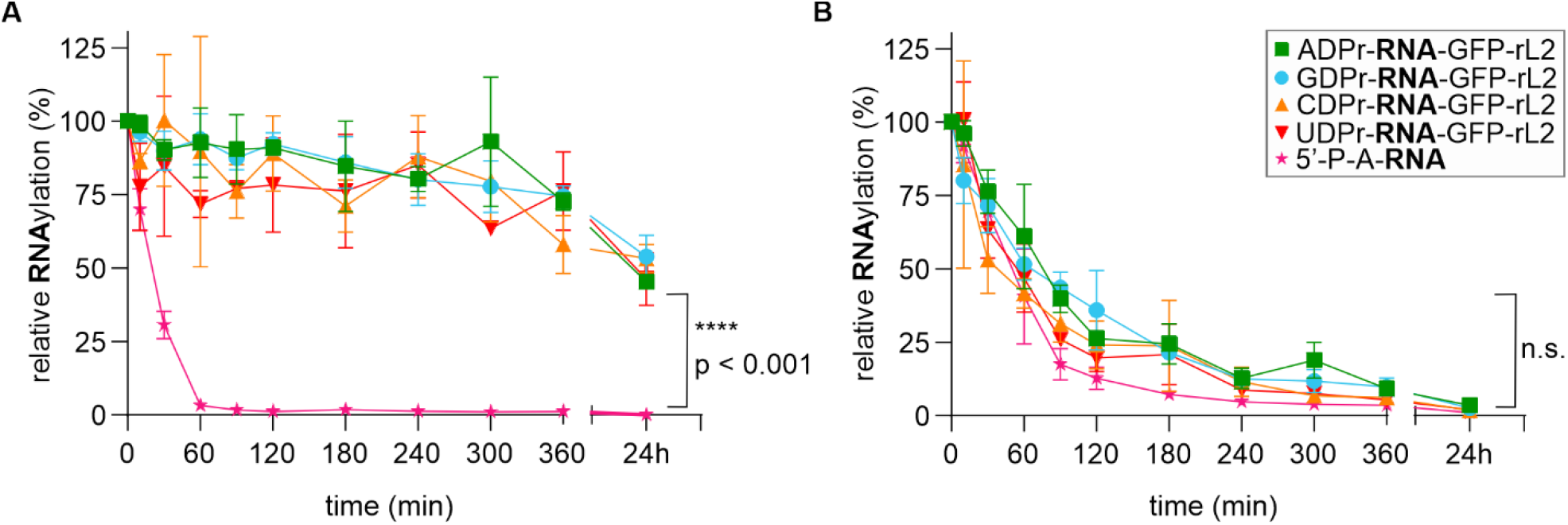
Cell lysate and blood plasma stability of NXD-RNAylated GFP-rL2. GFP-rL2 is RNAylated with 3’ Cy5 labelled NXD-RNA 10mers. A) The stability graph of NXD-RNAylated GFP-rL2 and control 5′-P-A-RNA in human embryonic kidney 293T (HEK293T) cell lysate is shown. Error bars represent the mean ± s.d. of n = 3 independent experiments. Statistical analysis is performed using one-way ANOVA with multiple comparisons in GraphPad Prism. Multiple comparisons are performed to assess statistical significance by comparing the relative RNAylation percentage of NXD-RNAylated GFP-rL2 with the relative detected abundance of 5’-P-A-RNA. Asterisks indicate significance levels: *p ≤ 0.05, **p ≤ 0.01, ***p ≤ 0.001, and ****p ≤ 0.0001. B) The stability graph of NXD-RNAylated GFP-rL2 and control 5′-P-A-RNA in mouse blood plasma is demonstrated. Error bars represent the mean ± s.d. of n = 3 independent experiments. Statistical analysis is performed using one-way ANOVA with multiple comparisons in GraphPad Prism. Multiple comparisons are performed to assess statistical significance by comparing the relative RNAylation percentage of NXD-RNAylated GFP-rL2 with the relative detected abundance of 5’-P-A-RNA. No statistical significance is observed.

We further assessed the stability of NXD-RNAylated GFP-rL2 in mouse blood plasma. As expected, 5′-P-A-RNA was highly labile, with 90% degraded after 2 hours of incubation. Surprisingly, NXD-RNAylated GFP-rL2 did not exhibit a stability advantage in blood plasma; it was degraded as efficiently as unmodified RNA (Figure 5B, Supplementary Figure 13A-E). This observation suggests that, while protein conjugation protects RNA from cytoplasmic exonucleases such as XRN1 and enhances stability in cell lysates, it does not confer resistance to the diverse and abundant nucleases present in blood plasma. Collectively, these findings demonstrate that covalent protein conjugation is an effective strategy to protect RNA from exonucleolytic degradation in cellular environments, but additional strategies may be required to achieve stability in extracellular fluids such as blood plasma.

### Purification and cellular delivery of DNAylated proteins

*In vitro* generated nucleic acid-protein conjugates can be applied for different purposes such as gene editing, biosensing, or therapeutic approaches. Their utility depends critically on achieving high purity to prevent off-target effects. To address this, we have established a two-step chromatographic purification protocol (Figure 6A). The main challenge in purifying these conjugates arises from the need to remove non-covalently associated nucleic acids, particularly since rS1 DII and other target proteins possess nucleic acid-binding domains. To ensure complete removal of residual nucleic acids, we have employed denaturing conditions during purification. The first step of the purification strategy is based on immobilized metal affinity chromatography (IMAC), where the His-tagged proteins are captured and residual nucleic acids are removed by urea washes (Figure 6A and Supplementary Figure 14A). This step is followed by anion exchange chromatography, which captures nucleic acid-protein conjugates based on ionic interactions. During this step, unmodified proteins and ModB are efficiently separated from the nucleic acid-protein conjugates, which are subsequently eluted in a highly purified form (Figure 6A and Supplementary Figure 14B).

**Figure 6:**
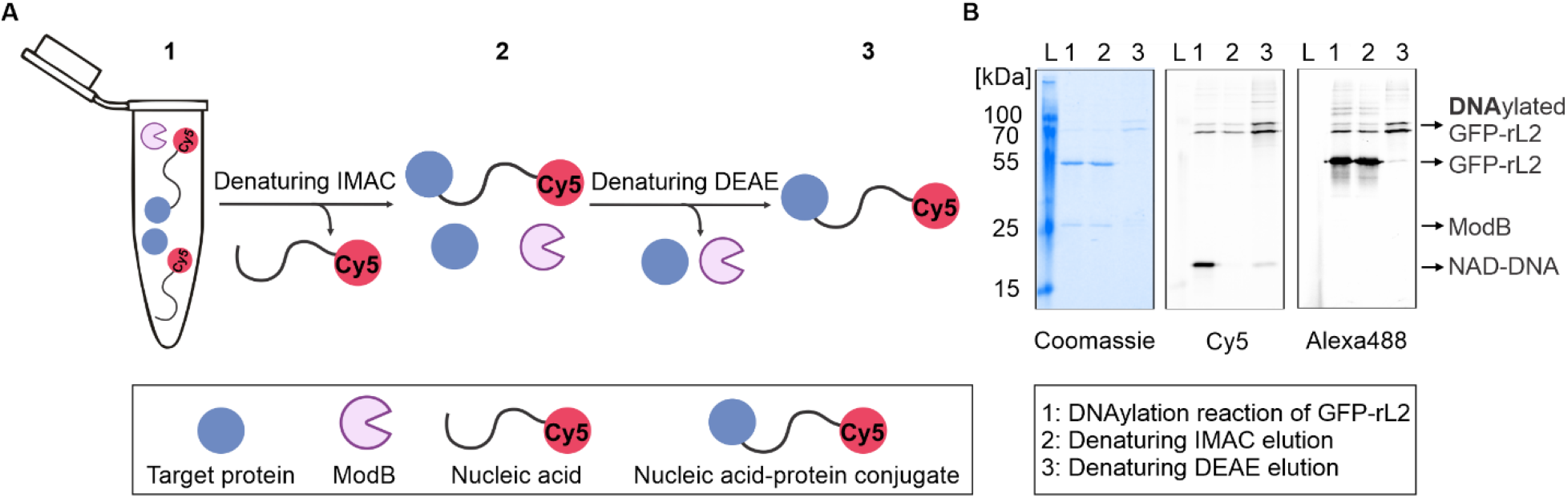
Established purification protocol for nucleic acid-protein conjugates. A) The purification protocol is explained schematically. The first purification step is denaturing immobilized metal affinity chromatography (IMAC) that enables the removal of non-covalently bound nucleic acids. The second purification is denaturing anion exchange chromatography performed with diethylaminoethyl (DEAE) resin. This step leads to the removal of the unmodified target protein and ModB. Upon a two-step purification protocol, a purified nucleic acid-protein conjugate is obtained. B) Purification products were analyzed by 12% SDS-PAGE.

To demonstrate the feasibility of delivering nucleic acid–protein conjugates into human cells, we have generated DNAylated GFP-rL2. Upon successful purification of DNAylated GFP-rL2 (Figure 6B), we have delivered purified DNAylated GFP-rL2 to HEK293T cells via liposome-based technologies. Confocal fluorescence microscopy confirmed the presence of the conjugate inside the transfected cells, with GFP fluorescence marking the protein and a 3′ Cy5-labeled NAD-DNA 60mer tracking the nucleic acid component (Figure 7). Co-localization of GFP and Cy5 signals in transfected cells provided clear evidence of successful intracellular delivery of DNAylated proteins (Figure 7). In contrast, control experiments using only protein or DNA showed signal exclusively in their respective channels (Supplementary Figure 15), confirming the specificity of the co-localization signal. These results establish that our purification strategy yields highly pure nucleic acid–protein conjugates, and that these conjugates can be efficiently delivered into human cells, confirming their promise for a range of biotechnological and therapeutic applications.

**Figure 7:**
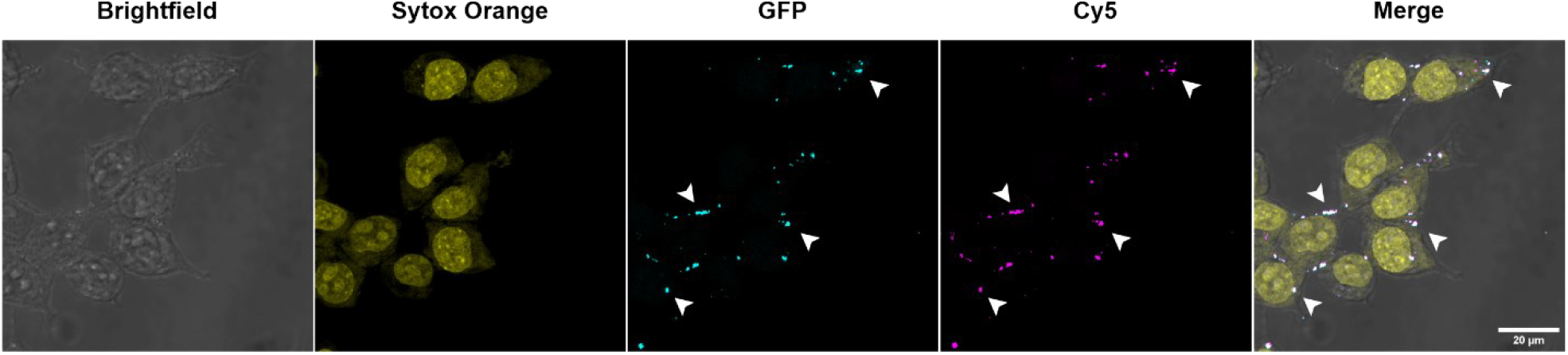
Maximum intensity projection images of DNAylated GFP-rL2 transfected HEK293T cells. Purified DNAylated GFP-rL2 delivered HEK293T cells are imaged via a fluorescence confocal microscopy. GFP-rL2 is DNAylated with 3’Cy5 labelled NAD-DNA 60mer. A maximum-intensity projection was generated from 20 consecutive z-sections (z-step: 0.37 microns), centered on the approximate mid-plane of the cell. Three fluorescence channels (Sytox Orange–stained nuclei, GFP, and Cy5) and the brightfield image are shown individually and as a merged composite. For clarity, selected co-localized intracellular signals are highlighted with arrowheads. (n = 3 independent experiments)

## Discussion

The development of site-specific, biocompatible nucleic acid–protein conjugates remains a central challenge in synthetic biology, molecular therapeutics, and nanotechnology. Here, we establish an enzymatic platform based on ModB-mediated RNAylation and DNAylation, enabling the covalent and site-selective attachment of diverse nucleic acids to proteins via a naturally occurring N-glycosidic bond. This approach addresses key limitations of existing chemical and biochemical conjugation strategies, which often lack site specificity, require labor-intensive protocols, and generate non-natural linkages that may compromise biocompatibility and trigger immunogenic responses *in vivo*^7, 8, 12, 14^-^17, 54^.

Our findings reveal that ModB displays remarkable and unexpected substrate promiscuity, efficiently catalyzing the conjugation of both NXD-capped RNAs and DNAs to target proteins. Strikingly, ModB tolerates the substitution of the adenine base in the NAD cap with guanine, cytosine, or uracil without any loss of activity, thereby greatly expanding the chemical diversity and design flexibility of nucleic acid–protein conjugates. This broad substrate tolerance enables the engineering of conjugates with precisely tailored resistance to enzymatic cleavage: while the N-glycosidic bond formed with ADPr-RNA is readily hydrolyzed by the eukaryotic enzyme ARH1^19^, conjugates generated with non-canonical caps (GDPr-, CDPr-, UDPr-RNA/DNA) exhibit pronounced stability against ARH1-mediated degradation. This property allows for the rational design of conjugates with tunable stability, supporting applications that demand either dynamic turnover or persistent functionality.

Furthermore, the broad substrate scope of ModB suggests that the enzyme has not been shaped by evolutionary selection pressure to discriminate among NXD-capped nucleic acid substrates, as only NAD-RNAs have been identified in nature^24, 25^. All other NXD-capped nucleic acids are synthetic and do not occur in biological systems, underscoring the unique versatility of ModB for biotechnological applications. Moreover, we observe four times higher conjugation yield in the presence of NCD-RNA compared to NAD-RNA. Surprisingly, among NXD-capped RNAs, the canonical substrate NAD-RNA yielded the lowest conjugate generation efficiency. On the other hand, NXD-capped DNAs lead to similar DNAylation efficiency, except NTD-DNA, where the efficiency was decreased by 40%. Given that no crystal structure of ModB is available yet, we cannot explain the observed trend for substrate preferences based on the predictions.

In this study, we establish a modular system that enables RNAylation of virtually any protein of interest *in vitro*. By defining an RNAylation tag composed of a fusion between a mutated version of SpyTag (R3K) and rS1 DII, we created a standardized tag that can be readily combined with any target protein, thereby ensuring site-selective modification. Notably, we deliberately introduced an arginine-to-lysine mutation within the SpyTag sequence to prevent unintended RNAylation at the SpyTag itself, which was observed when the native arginine was present. This strategic mutation confines RNAylation to the designated arginine residue within rS1 DII, enabling precise and predictable site-specific conjugation. Importantly, the introduction of the R3K mutation does not impair the efficiency of the SpyCatcher–SpyTag reaction, as the N-terminal arginine is not essential for covalent bond formation between SpyTag and SpyCatcher^38^. Such control over the modification site is essential for pharmaceutical and biotechnological applications, where guaranteeing RNAylation at a specific residue is critical for the function, safety, and reproducibility of nucleic acid-protein conjugates.

A key advantage of ModB-mediated conjugation is the formation of a natural, biologically recognized N-glycosidic bond^19^, which contrasts with the non-cleavable thioether bonds produced by widely used tags such as SNAP-tag^16, 17^. The natural linkage not only enhances biocompatibility but also allows for regulated degradation and recycling of conjugates by endogenous enzymes, aligning with cellular processes and reducing the risk of long-term immunogenicity.

Furthermore, the data reveal that covalent attachment of RNA to proteins markedly enhances RNA stability in human cell lysates, protecting against exonucleolytic degradation and extending the functional lifetime of RNA in cellular environments. However, this protective effect does not extend to blood plasma, where abundant extracellular nucleases and proteases rapidly degrade both free and conjugated RNA^55-57^. These findings highlight the need for additional strategies-such as protein engineering^58-61^, RNA modifications^62-65^, or encapsulation^66^ to improve the pharmacokinetic properties and extracellular stability of nucleic acid–protein conjugates for therapeutic applications. To overcome this challenge, current RNA therapeutics employ protective delivery strategies to prevent RNA degradation in blood plasma. Lipid nanoparticles (LNPs) have emerged as the most widely used platform, offering efficient encapsulation, enhanced circulation time, and improved cellular uptake^67^.

This study also provides a purification strategy for RNAylated-proteins. The utilization of a pure product is essential for many applications, such as therapeutic approaches or biotechnological applications^68^. Therefore, we have developed an efficient purification strategy to obtain pure nucleic acid-protein conjugates. After successfully generating purified DNA-protein conjugates, we demonstrated their delivery into HEK293T cells as a proof of concept. This validates that DNAylated proteins can be introduced into cells using conventional delivery methods, establishing their potential for broader applications in intracellular targeting and functional studies.

Although our conjugation platform presents a powerful strategy to produce nucleic acid-protein conjugates, it has certain limitations that must be addressed for broader applicability. While we have established design principles, evaluated conjugate stability under physiological conditions, developed an efficient purification workflow, and successfully delivered the purified conjugates to human cells, the functional potential of conjugates remains to be validated in biological systems. Another limitation of the study is that, despite being smaller than conventional fusion tags such as SNAP-tag (20 kDa)^16^, rS1 DII (10.7 kDa)^37^ must still be fused to SpyTag or to a POI directly to make the attachment of nucleic acids possible, which may impact protein structure or function. Future efforts should focus on optimizing and truncating rS1 DII to further minimize its size while retaining or enhancing its efficiency, reducing potential steric hindrance. Addressing these limitations through further engineering and refinement will be crucial for unlocking the full potential of ModB-mediated conjugation as a versatile tool. To conclude, in this study, we provide a foundational framework for generating site-specific nucleic acid-protein conjugates facilitated by ModB in a single-step enzymatic reaction, establishing a basis for its broader application to generate functional conjugates to be applied in the synthetic biology or therapeutics field.

## Author contributions

Nurseda Yilmaz Demirel: Conceptualization, Formal analysis, Investigation, Methodology, Validation, Writing—original draft. Ilayda Mumcuoglu: Investigation, Validation. Gabriele Malengo: Investigation, Methodology. Bernd Schmeck: Conceptualization. Anna Lena Jung: Conceptualization, Methodology. Katharina Höfer: Conceptualization, Methodology, Writing-original draft and review & editing, Funding acquisition.

## Acknowledgements

We would like to thank Jun.-Prof. Dr. rer. nat. Leon Schulte for generously sharing the HEK293T cell line, and to Prof. Dr. Oliver Hantschel for the mouse blood plasma. Additionally, we would like to thank Dr. Timo Glatter for LC-MS/MS sample analysis.

## Funding

This work has received funding from the European Research Council under the European Union’s Horizon 2020 research and innovation programme (ERC-2023-STG grant 101114948, NAD-ART to K.H.), from the Max Planck Society (Max Planck Research Group Leader funding to K.H.), the DFG-funded research consortium GRK 2937 (project number 505997786 to K.H.), the Hessisches Ministerium für Wissenschaft und Kunst (LOEWE Diffusible Signals LOEWE-Schwerpunkt Diffusible Signals to B.S. and A.L.J.; LOEWE Exploration to B.S.), and the von Behring Roentgen foundation (to A.L.J.).

## Declaration of competing interest

K.H. is in the process of applying for a patent (PCT/EP2021/071295) covering the RNAylation that lists K.H. as inventor. The remaining authors declare that they have no known competing financial interests or personal relationships that could have appeared to influence the work reported in this paper.

## Methods

### General methods

3’ Cy5-labeled 5’-monophosphorylated-RNA and 5’-monophosphorylated-DNA oligos (Supplementary Table 1) were purchased as HPLC-purified oligos from Integrated DNA Technologies (IDT). ChemiDoc MP from BioRad was used to visualize SDS and Tricine gels, and Image Lab software from BioRad was used for image analysis. Coomassie staining was performed with Instant Blue Coomassie stain purchased from Sigma-Aldrich. Glycerol, imidazole, tricine, xylene cyanol, Rotiphorese NF-Acrylamide/Bis-solution 30%, Rotiphorese sequencing gel concentrate, and Rotiphorese sequencing gel diluter were purchased from Carl Roth. EDTA (Ethylenediamine tetraacetic acid) and Bromophenol blue were purchased from Serva. Urea was supplied by Labochem International. Additional reagents were purchased from Sigma-Aldrich and used without further purification. Protein concentration was measured with the NanoDrop ND-1000 Spectrophotometer (Thermo Fisher Scientific).

### AlphaFold3 prediction

The structural prediction of ModB is performed via AlphaFold3^47^ in default settings with a single copy of ModB (Uniprot ID: P39423, MODB_BPT4) protein and a single copy of the NAD substrate. Among generated predictions, the one with the highest predicted Template Modeling (pTM) value of 0.89, showing high overall confidence in the global structure arrangement, has been taken for further analysis. The color-coding of per-residue confidence, which is evaluated by using predicted Local Distance Difference Test (pLDDT) scores, is visualized in PyMol by using the color code from AlphaFold. The pLDDT color scheming for the structural visualization is as follows: blue (≥90, very high confidence), cyan (70–89, confident), yellow (50–69, low confidence), and orange to red (<50, very low confidence).

### Cloning of target proteins

*E. coli* One Shot BL21 (DE3) strains of *pET28a*-ModB-His_6_, *pET28a*-DII-His_6_, *pET28c*-rS1-His_6,_ and *pET28c*-rL2-His_6_ were produced in Wolfram-Schauerte *et al*.^19^ *pJ404*-SpyCatcher003-sfGFP was purchased from Addgene (catalog number 133449). SpyTag003 sequence (MGRGVPHIVMVDAYKRYK) was introduced to *pET28a*-DII-His_6_ with the primers listed in the Supplementary Table 2 via PCR, followed by self-circularization, resulting in *pET28a*-SpyTag-DII-His_6_. To generate SpyTag R3K-DII version, single point mutation was introduced to the *pET28a*-SpyTag-DII-His_6_ plasmid with the primers sequences are listed in Supplementary Table 2. sfGFP sequence is introduced (the inserted sequence is written in the Supplementary Table 2) to *pET28a*-rL2-His_6_ plasmid by Gibson Assembly (NEB Gibson Assembly Master Mix, E2611S) with the primers written in the Supplementary Table 2. *pET28c*-rL2-His_6_ was used to amplify the backbone for the Gibson Assembly and *pJ404*-SpyCatcher003-sfGFP was used to amplify sfGFP insert sequence.

### Protein purification

Except for ModB, all purified proteins utilized for *in vitro* studies are overexpressed and purified in the same way.

*E. coli* One Shot BL21 (DE3) containing the C-terminal His-tagged (His_6_) respective protein in pET28a plasmid was cultured in LB medium at 37 °C. Protein expression was induced with isopropyl β-d-1-thiogalactopyranoside (IPTG) at OD_600_ = 0.8 for 3 hours at 37 °C. Bacteria were harvested by centrifugation (3,800 x g, 4 °C, 15 min) and lysed by sonication (5 min, 50 % power, two times) in HisTrap buffer A (50 mM Tris-HCl pH 7.5, 1 M NaCl, 1 M Urea, 5 mM MgSO4, 5 mM ß-mercaptoethanol, 5 % glycerol, 5 mM imidazole, 1 tablet per 500 ml complete EDTA-free protease inhibitor cocktail (Roche)). The lysate was clarified by centrifugation (37,500 x g, 30 min, 4 °C), and the supernatant was sterile-filtered and then applied to a 1 mL Ni-NTA HisTrap column (Cytiva). The protein was eluted with an imidazole gradient using HisTrap buffer B (HisTrap buffer A with 300 mM imidazole) and analyzed by 12% SDS–PAGE or 15% Tricine-PAGE.

Further protein purification was achieved by size exclusion chromatography (SEC) through a Superdex 200 300/10 GL column (Cytiva) using size exclusion buffer containing 300 mM NaCl and 50 mM Tris-HCl, pH 7.5. Fractions of interest were analyzed by 12% SDS–PAGE or 15% Tricine-PAGE, pooled and concentrated in Amicon Ultra-4 centrifugal filters (MWCO 3 kDa or 10 kDa, centrifugation at 4,000 x g, 4 °C). Proteins were finally stored in buffer supplemented with 50 % glycerol at −20 °C.

### Purification of ModB protein

*E. coli* One Shot BL21 (DE3) *pET28a*-ModB-His_6_ was grown to an OD_600_ = 1.2 at 37 °C, 180 rpm, and cooled down to 4 °C while shaking at 160 rpm for at least 30 min. The addition of 1 mM IPTG induced protein expression. The cultures were then incubated for 120 min at 4 °C, 160 rpm, and bacteria were harvested by centrifugation (3,800 x g, 4 °C, 15 min). The rest of the protocol is the same as the protein purification protocol explained above, starting from the sonication step.

### Preparation of 5’ NXD-capped nucleic acids

5’-monophosphorylated-RNAs and 5’-monophosphorylated-DNAs (sequences can be found in Table 1) were subjected to an imidazolide reaction to generate 5’ NXD-capped RNAs and 5’ NXD-capped DNAs, respectively^49^. 5’-monophosphorylated oligos were incubated with a 1000-fold excess of nicotinamide mononucleotide (NMN)-phosphorimidazolide (Im-NMN) (synthesized based on the published protocol^49^), 50 mM MgCl_2_ for 5 h at 50 °C. The remaining Im-NMN in the reaction was eliminated using Amicon Ultra 3 kDa filters. The reaction mixtures were eluted from the filter to fresh tubes. Capping efficiencies were analyzed by analytical APB (acryloyl aminophenyl boronic acid) gel electrophoresis^69^. APB gel was prepared by dissolving 30 mg of APB in 2 mL 10x HAE Buffer (200 mM HEPES-NaOH, pH 8.2, 50 mM NaOAc, 10 mM EDTA), 10 mL Rotiphorese sequencing gel concentrate, and 8 mL Rotiphorese sequencing gel diluter. The mixture was heated to 50 °C for 5 min. After APB was dissolved, the mixture was cooled down before adding 100 µL 10% ammonium persulfate (APS) and 15 µL N, N, N’, N’ Tetramethylethylenediamine (TEMED).

Imidazolide reaction samples and 5’-P-X-RNAs as negative controls were run on an APB gel at 5W for 30 min to analyze 5’ NXD-capping. The band intensities of imidazolide reaction samples were analyzed using the Image Lab software. By accepting the total band intensity of a 5’ NXD-nucleic acid and a 5’-P-X-nucleic acid as 100%, the capping efficiency of the imidazolide reaction was calculated for each Im-NMN reaction.

### RNAylation and DNAylation of proteins *in vitro*

For RNAylation and DNAylation reactions, 2 pmol/µL of each 5’ NXD-capped nucleic acid was incubated with 1 µM ModB and 2 µM target protein in the presence of 1x Transferase buffer (10 mM Mg(OAc)_2_, 22 mM NH_4_Cl, 50 mM Tris-acetate pH 7.5, 1 mM EDTA, 10 mM ß-mercaptoethanol, 1% glycerol) at 15 °C for 2.5 hours if another recipe is not stated in the methods section. The reaction was stopped by adding 2x Laemmli loading dye (100 mM Tris-HCl pH 6.8, 4% SDS, 20% glycerol, 2% β-mercaptoethanol, 6.25 mM EDTA, and a tip of spatula of Bromophenol blue) and analyzed by 12% SDS-PAGE. Cy5 signal corresponding to the reaction was visualized using ChemiDoc MP. Following Cy5 imaging, gels were stained with Coomassie Blue overnight and destained with water. Coomassie staining was imaged with ChemiDoc MP. Image analysis was performed in Image Lab software. The Cy5 and Coomassie staining band intensities were calculated using Image Lab software to quantify RNAylation reaction efficiencies. Cy5 band intensity of 5’ NXD-capped nucleic acid reaction in the presence of ModB was normalized to the respective Coomassie band intensity. These calculated ratios were normalized to the RNAylation efficiency of the sample 5’ NAD-capped nucleic acid in the presence of ModB, which is used as a reference.

### LC-MS/MS analysis of RNAylated SpyTag-DII

The sample preparation for the RNAylated residues on SpyTag-DII is performed according to the published methods in Wolfram-Schauerte *et al*.^19^. *In vitro* RNAylated SpyTag-DII was directly digested (without further purification) with 1 μg RNase A (Thermo Fisher Scientific) and 100 U RNase T1 (Thermo Fisher Scientific) at 37 °C for 1 h, following tryptic digestion at 37 °C for 3 h in the same buffer with trypsin (Promega) in a 1:30 ratio (w/w) relative to the total protein content in each sample. Peptides were purified with Chromabond C18 WP spin columns and submitted for LC–MS/MS analysis. The sample run and data processing were performed by Dr. Timo Glatter with the same settings for sample run and data analysis as published in Wolfram-Schauerte *et al*.^19^

### Modular system establishment

Modular system establishment was carried out at 25 °C in succinate–phosphate–glycine buffer (12.5 mM succinic acid, 43.75 mM NaH_2_PO_4_, 43.75 mM glycine; pH adjusted to 7.0 using NaOH)^70^ for 45 min. To test modular system establishment with SpyTag-DII, 2 µM SpyCatcher-GFP was incubated with either 4 µM SpyTag-DII or RNAylated SpyTag-DII. To establish a modular system in the presence of SpyTag R3K-DII, 20 µM SpyCatcher-GFP was incubated with either 40 µM SpyTag R3K-DII or RNAylated SpyTag R3K-DII. RNAylation reactions were performed according to the protocol written in the RNAylation and DNAylation of proteins *in vitro* section. The reaction was stopped by adding 2x Laemmli loading dye and analyzed by 12% SDS-PAGE.

### ARH1 kinetics of RNAylated and DNAylated rS1

RNAylation and DNAylation reactions of rS1 have been performed as explained in the RNAylation and DNAylation of proteins *in vitro* section. 1x Transferase buffer was exchanged to 1x Degradation Buffer (12.5 mM Tris-HCl pH 7.5, 25 mM NaCl, 25 mM KCl, 5 mM MgCl_2_, 1 mM β-mercaptoethanol) using Zeba Spin Desalting Columns (7K MWCO, Thermo Fisher Scientific). 0.1 µM RNAylated rS1 was digested with 0.004 µM ARH1 (Enzo, 0.2 mg/mL) at 37 °C for 0, 2, 5, 10, 30, 60,120, and 180 minutes. Digestion reactions were stopped by adding 2x Laemmli loading dye and analyzed by 12% SDS-PAGE. Cy5 signaling was visualized by ChemiDoc MP. Following Cy5 imaging, gels were stained with Coomassie Blue overnight and destained with water. Coomassie staining was imaged with ChemiDoc MP. Image analysis was performed in Image Lab software. The Cy5 and Coomassie staining band intensities were calculated using Image Lab software to quantify relative RNAylation percentages. Cy5 band intensities were normalized to respective Coomassie staining intensities. 5’ NXD-capped nucleic acid 0 min reaction was accepted as 100%, and the calculated ratios were normalized to the respective 5’ NXD-capped nucleic acid 0 min reactions. Final calculations were transferred to OriginPro software. Means and standard deviations were calculated for the two data sets and plotted as graphs in the software. Non-linear fit curves of data points were shown.

### Xrn1 kinetics of RNAylated rS1

RNAylation reactions of rS1 have been performed as explained in the RNAylation and DNAylation of proteins *in vitro* section. 1x Transferase buffer was exchanged to 1x Degradation Buffer (12.5 mM Tris-HCl pH 7.5, 25 mM NaCl, 25 mM KCl, 5 mM MgCl_2_, 1 mM β-mercaptoethanol) using Zeba Spin Desalting Columns (7K MWCO, Thermo Fisher Scientific). 0.1 µM RNAylated rS1 was digested with 0.005 U/µL XRN-1 (NEB,1 U/µL) at 37 °C for 0, 2, 5, 10, 30, 60,120, and 180 minutes. Digestion reactions were stopped by adding 2x Laemmli loading dye and analyzed by 12% SDS-PAGE. Cy5 signaling was visualized by ChemiDoc MP. Following Cy5 imaging, gels were stained with Coomassie Blue overnight and destained with water. Coomassie staining was imaged with ChemiDoc MP. Image analysis was performed in Image Lab software. The Cy5 and Coomassie staining band intensities were calculated using Image Lab software to quantify relative RNAylation percentages. Cy5 band intensities were normalized to respective Coomassie staining intensities. 5’ NXD-capped nucleic acid 0 min reaction was accepted as 100%, and the calculated ratios were normalized to the respective 5’ NXD-capped nucleic acid 0 min reactions. Final calculations were transferred to OriginPro software. Means and standard deviations were calculated for the two data sets and plotted as graphs in the software. Non-linear fit curves of data points were shown.

### Nucleic acid-protein conjugates purification strategy

Purification of nucleic acid-protein conjugates is based on a two-step chromatographic purification protocol. The first step is denaturing immobilized metal affinity chromatography (IMAC), which aims to remove unattached nucleic acids. For this purpose, Ni-NTA magnetic beads (Serva, 42179.02) were used to capture His-tagged proteins. The bead amount is determined based on the total protein amount applied for the purification (Please check the manufacturer’s guidelines to determine the required bead volume). Ni-NTA magnetic beads were washed three times with Wash Buffer 1 (50 mM Tris-HCl pH=7.5, 20 mM Imidazole, 1 M NaCl, 1 M Urea, 5 mM MgSO_4_, 5 mM β-mercaptoethanol, 5% Glycerol, EDTA-free protease inhibitor) supplemented with 0.1% Tween-20. Afterward, the beads were equilibrated three times with Wash Buffer 1. After bead equilibration, a sample is prepared by adding 1 mL of Wash Buffer 1 to 300 µL DNAylated GFP-rL2 reaction (containing approximately 150 µg total protein), and the mixture is applied to 300 µL equilibrated beads. The sample has been incubated at 4°C for 1 hour on a rotary. After incubation, the flow-through was collected. The beads were washed four times with Wash Buffer 1, where the second wash step is supplemented with 0.1% Tween-20 to maintain bead solubility. To remove non-covalently interacting attached nucleic acids, three times Urea Buffer (50 mM Tris-HCl pH=7.4, 8M Urea) wash was performed. Following, the beads have been washed five times with Wash Buffer 1. The elution was performed with Elution Buffer 1 (50 mM Tris-HCl pH=7.5, 500 mM Imidazole, 500 mM NaCl, 1 M Urea, 5 mM MgSO_4_, 5 mM β-mercaptoethanol, 5% Glycerol, 0.1% Tween-20, EDTA-free protease inhibitor). Based on the bead volume, the elution volume needs to be decided. 250 µL Elution Buffer 1 was used based on the manufacturer’s guidelines. The purity of the elution product was analyzed via 12%SDS-PAGE.

The elution product from the first step is diluted 24 times the volume of elution with Wash Buffer 2 (50 mM Tris-HCl pH=7.5, 8M Urea) to enable diethylaminoethyl (DEAE) column binding (Vivapure D Mini, Sartorius, VS-IX01DH24). After the spin column is equilibrated with Wash Buffer 2, the sample is applied. Upon sample application, the column is washed 10 times with Wash Buffer 2. This is followed by 5 times wash with Wash Buffer 2 + 0.2 M NaCl to remove nonspecifically interacting biomolecules. Elution is performed with Elution Buffer 2 (50 mM Tris-HCl pH=7.5, 8M Urea, 1 M NaCl). Elution has been performed two times with 100 µL Elution Buffer 2 to concentrate the sample. The purity of the sample was analyzed via 12% SDS-PAGE.

To achieve buffer exchange of a DEAE elution sample with a small volume (approximately 100 µL), dialysis filters (V-series, 0.025 µm pore size with a filter diameter of 47 mm, Merck, VSWP04700) are utilized. For dialysis, a container is filled with 30 mL of 50 mM Tris-HCl, pH 7.5. The dialysis filter has been placed in the middle of the container with a blunt-end forcep (Merck, XX6200006P) very carefully. 100 µL of elution sample has been applied to the center of the filter. The container has been closed with a lid to prevent evaporation. The dialysis has been performed at 4°C for 30 minutes. Upon incubation, the dialysed sample is transferred into a new Eppendorf tube and stored at −20°C for further use.

### Cell lysate generation and stability assay

Human embryonic kidney (HEK) 293T cell lysate was prepared under non-denaturing conditions to maintain cellular enzymatic activity. HEK293T cells were grown in a T175 flask to 80-90% confluency in DMEM (Dulbecco’s Modified Eagle Medium) (Fisher Scientific, 11965092) + 10% FBS (Fetal Bovine Serum) (Fisher Scientific, A5670701). Cells were washed twice with ice-cold Gibco PBS (phosphate buffered saline), pH=7.4, without calcium and magnesium (Fisher Scientific, 10010023). Cells were scraped in 10 mL of non-denaturing lysis buffer (50 mM Tris-HCl, 150 mM NaCl, 0.5% Triton X-100, pH=7.5). To complete the cell lysis, cells were sonicated 5 times at 20% amplitude for 5 seconds. During sonication, cells were incubated on ice, and between each sonication step, a 30-second break was taken to allow the sample to cool down. After sonication, the sample was centrifuged at 13,000 x g for 10 minutes at 4°C to remove cellular debris. Supernatant has been collected and aliquoted. The aliquots were flash frozen to be stored at −80°C until further use. One aliquot is used to determine the protein concentration of the generated lysate with the BCA assay (Pierce BCA Protein Assay Kit, Scientific Fisher, 23250).

NXD-RNAylated GFP-rL2 protein was produced in the presence of 1 µM NXD-RNA 10mer Cy5, 5 µM GFP-rL2, and 5 µM ModB in 1x Transferase buffer (10 mM Mg(OAc)_2_, 22 mM NH_4_Cl, 50 mM Tris-acetate pH=7.5, 1 mM EDTA, 10 mM ß-mercaptoethanol, 1% glycerol) at 15°C for 2.5 hours. To remove non-attached NXD-RNA from the samples, denaturing IMAC purification is performed as explained above. Elution products are buffer exchanged 1x Degradation Buffer (12.5 mM Tris-HCl pH=7.5, 25 mM NaCl, 25 mM KCl, 5 mM MgCl_2_, 1 mM ß-mercaptoethanol) using Zeba Spin Desalting Columns (7K MWCO, Thermo Fisher Scientific).

1 µM of NXD-RNAylated rS1 and 2 µM of 5’-P-A-RNA 10mer Cy5 were incubated with cell lysate (final concentration: 0.7 mg/mL) at 37 °C for 0, 10, 30, 60,120, 180, 240, 300, 360 minutes, and 24 hours. Digestion reactions were stopped by adding 2x Laemmli loading dye and analyzed by 12% SDS-PAGE. Cy5 signaling and Stain-free intensities were visualized by ChemiDoc MP. Image analysis was performed in Image Lab software. The Cy5 and total lane Stain-Free band intensities were calculated using Image Lab software to quantify relative RNAylation percentages. Cy5 band intensities were normalized to respective total lane Stain-Free intensities. 5’ NXD-capped nucleic acid 0 min reaction was accepted as 100%, and the calculated ratios were normalized to the respective 5’ NXD-capped nucleic acid 0 min reactions. Final calculations were transferred to OriginPro software. Means and standard deviations were calculated for the two data sets and plotted as graphs in the software. Non-linear fit curves of data points were shown. Following Cy5 imaging, gels were stained with Coomassie Blue overnight and destained with water. To assess the 5’-P-A-RNA 10mer Cy5 stability in cell lysate, samples were collected at the same time points and mixed with colorless 2x PAGE loading dye (90% Formamide+10% 10x TBE (1M Tris, 1M Boric acid, 25 mM EDTA) to stop the reaction. The samples were analyzed via 20% PAGE. Cy5 signaling was visualized by ChemiDoc MP. Image analysis was performed in Image Lab software. The Cy5 band intensities were calculated using Image Lab software to quantify relative RNA amount. 5’-P-A-RNA nucleic acid 0 min reaction was accepted as 100%, and the calculated ratios were normalized to it. Final calculations were transferred to OriginPro software. Means and standard deviations were calculated for the two data sets and plotted as graphs in the software. Non-linear fit curves of data points were shown.

### Blood plasma stability assay

NXD-RNAylated GFP-rL2 protein was produced in the presence of 1 µM NXD-RNA 10mer Cy5, 5 µM GFP-rL2, and 5 µM ModB in 1x Transferase buffer (10 mM Mg(OAc)_2_, 22 mM NH_4_Cl, 50 mM Tris-acetate pH=7.5, 1 mM EDTA, 10 mM β-mercaptoethanol, 1% glycerol) at 15°C for 2.5 hours. To remove non-attached NXD-RNA from the samples, denaturing IMAC purification is performed as explained above. Elution products are buffer exchanged 1x Degradation Buffer (12.5 mM Tris-HCl pH=7.5, 25 mM NaCl, 25 mM KCl, 5 mM MgCl_2_, 1 mM ß-mercaptoethanol) using Zeba Spin Desalting Columns (7K MWCO, Thermo Fisher Scientific).

1 µM of NXD-RNAylated rS1 and 2 µM of 5’ P-A-RNA 10mer Cy5 were incubated with mouse blood plasma (Maus Plasma Non-Sterile in Sodium Heprain, Antikoerper, ABIN925342) (final concentration: 0.8 mg/mL) at 37 °C for 0, 10, 30, 60,120, 180, 240, 300, 360 minutes, and 24 hours. Digestion reactions were stopped by adding 2x Laemmli loading dye and analyzed by 12% SDS-PAGE. Cy5 signaling and Stain-free intensities were visualized by ChemiDoc MP. Image analysis was performed in Image Lab software. The Cy5 and total lane Stain-Free band intensities were calculated using Image Lab software to quantify relative RNAylation percentages. Cy5 band intensities were normalized to respective total lane Stain-Free intensities. 5’ NXD-capped nucleic acid 0 min reaction was accepted as 100%, and the calculated ratios were normalized to the respective 5’ NXD-capped nucleic acid 0 min reactions. Final calculations were transferred to OriginPro software. Means and standard deviations were calculated for the two data sets and plotted as graphs in the software. Non-linear fit curves of data points were shown. Following Cy5 imaging, gels were stained with Coomassie Blue overnight and destained with water. To assess the 5’ P-A-RNA 10mer Cy5 stability in cell lysate, samples were collected at the same time and mixed with colorless 2x PAGE loading dye (90% Formamide+10% 10x TBE (1M Tris, 1M Boric acid, 25 mM EDTA) to stop the reaction. The samples were analyzed via 20% PAGE. Cy5 signaling was visualized by ChemiDoc MP. Image analysis was performed in Image Lab software. The Cy5 band intensities were calculated using Image Lab software to quantify relative RNA amount. 5’ P-A-RNA nucleic acid 0 min reaction was accepted as 100%, and the calculated ratios were normalized to it. Final calculations were transferred to OriginPro software. Means and standard deviations were calculated for the two data sets and plotted as graphs in the software. Non-linear fit curves of data points were shown.

### Cell delivery of DNAylated GFP-rL2

GFP-rL2 protein was DNAylated in the presence of 1 µM NXD-DNA 60mer Cy5, 5 µM GFP-rL2, and 5 µM ModB in 1x Transferase buffer (10 mM Mg(OAc)_2_, 22 mM NH_4_Cl, 50 mM Tris-acetate pH=7.5, 1 mM EDTA, 10 mM β-mercaptoethanol, 1% glycerol) at 15°C for 2.5 hours. The DNAylation product was purified as explained in the nucleic acid-protein conjugates purification strategy section. Upon confirming the purity via 12% SDS-PAGE, dilutions of 5’-P-X-DNA 60mer Cy5 and GFP-rL2 with the purified DNAylated GFP-rL2 have been analyzed via 12% SDS-PAGE to perform gel-based quantification. The band intensities are analyzed in ImageLab software. A linear graph for each fluorophore was generated based on the calculated band intensities in Origin software; these graphs have been used for interpolation of DNAylated GFP-rL2 to calculate the concentrations of each fluorophore for deciding the experimental control conditions. For transfection, 2 × 10^4^ HEK293T cells have been seeded in 300 µL DMEM+10% FBS per well of µ-slide 8 well ibiTreat (80826, 8-well glass bottomed). Seeded cells were incubated overnight at 37 °C 5% CO_2_. The next morning, the medium was replaced with 250 µL DMEM. 25 µL OptiMEM (Fischer Scientific) and 0.75 µL Lipofectamine MessengerMax (Fischer Scientific) are mixed and incubated at room temperature for 10 minutes. In the meantime, 15 µL of each sample is mixed with 10 µL OptiMEM. When 10 minutes of incubation is done, 25 µL Lipofectamine MessengerMax mixture and 25 µL sample were mixed and incubated at room temperature for 5 minutes. Upon incubation, the mixture is added to the respective wells dropwise and incubated at 37 °C 5% CO_2_ for 6 hours. After 6 hours of incubation, cell fixation and staining steps were performed. The medium was aspirated, and cells were washed three times with 1x PBS to remove residual culture medium. Under a chemical fume hood, 300 µL of 4% PFA in PBS was added to each well and incubated at room temperature for 10 minutes. After carefully removing the 4% PFA, cells were washed three times with 1x PBS. To lyse the cells, 300 µL of 0.1% Triton X-100 in PBS was added to the fixed cells. Cells were incubated for 5 minutes at room temperature. After carefully removing the 0.1% Triton X-100, cells were washed three times with 1x PBS. At this stage, cells are stored in the dark at 4°C overnight. The next morning, 1x PBS was removed, and 300 µL of freshly prepared 2 µM Sytox Orange working solution was added to the cells. The cells were incubated for 20 minutes at 4°C in the dark. After removing the staining solution, cells were washed three times with 1x PBS. Following this step, 1x PBS is exchanged with FluoroBright DMEM for confocal microscopy imaging.

### Confocal microscopy imaging and analysis

HEK293T fixed cells were imaged using an LSM 880 Zeiss Confocal microscope equipped with a Plan-Apochromat 63x/1.40 NA objective; the pinhole size was set to 0.93 Airy Units. Three excitation laser lines were reflected simultaneously using a 488/543/633 nm beam splitter. Three fluorescence channels were detected: GFP (Excitation: 488 nm; Emission: 490-544 nm); Sytox Orange (Excitation 543 nm; Emission 570-588 nm); Cy5 (Excitation 633 nm; Emission 635-735 nm). Moreover, brightfield images were acquired using the T-PMT. Z-stacks were acquired at 0.37 μm intervals, over a total depth of approximately 15-20 μm, corresponding to the typical cellular thickness. Images were processed using Fiji (ImageJ). For each z-stack, the Stack Focuser plugin was applied to the brightfield channel to identify the most in-focus plane. Based on this reference, a subset of 21 planes (±10 around the identified focal plane) was selected for maximum-intensity projection. Merged images were created using Fiji.

### Statistical analysis

Statistical analyses were performed in Origin 2020 and GraphPad Prism 10.

## References

1. Kulkarni, J.A. et al. The current landscape of nucleic acid therapeutics. Nature Nanotechnology 16, 630–643 (2021).

2. Moccia, M. et al. Advances in Nucleic Acid Research: Exploring the Potential of Oligonucleotides for Therapeutic Applications and Biological Studies. International Journal of Molecular Sciences 25, 146 (2024).

3. Stephanopoulos, N. Hybrid Nanostructures from the Self-Assembly of Proteins and DNA. Chem 6, 364–405 (2020).

4. Ramakrishnan, V. & White, S.W. Ribosomal protein structures: insights into the architecture, machinery and evolution of the ribosome. Trends in biochemical sciences 23, 208–212 (1998).

5. Corley, M., Burns, M.C. & Yeo, G.W. How RNA-binding proteins interact with RNA: molecules and mechanisms. Molecular cell 78, 9–29 (2020).

6. Adler, M. in Oxford Handbook of Nanoscience and Technology: Volume 2: Materials: Structures, Properties and Characterization Techniques. (eds. A.V. Narlikar & Y.Y. Fu) 0 (Oxford University Press, 2010).

7. Kashyap, D. & Booth, M.J. Nucleic Acid Conjugates: Unlocking Therapeutic Potential. ACS Bio & Med Chem Au 5, 3–15 (2025).

8. Dugal-Tessier, J., Thirumalairajan, S. & Jain, N. Antibody-Oligonucleotide Conjugates: A Twist to Antibody-Drug Conjugates. Journal of Clinical Medicine 10, 838 (2021).

9. Osada, E. et al. Engineering RNA–Protein Complexes with Different Shapes for Imaging and Therapeutic Applications. ACS Nano 8, 8130–8140 (2014).

10. Shibata, T. et al. Protein-driven RNA nanostructured devices that function in vitro and control mammalian cell fate. Nature Communications 8, 540 (2017).

11. Teodori, L., Omer, M. & Kjems, J. RNA nanostructures for targeted drug delivery and imaging. RNA Biol 21, 1–19 (2024).

12. Gokulu, I.S. & Banta, S. Biotechnology applications of proteins functionalized with DNA oligonucleotides. Trends in Biotechnology 41, 575–585 (2023).

13. Hoyt, E.A., Cal, P.M.S.D., Oliveira, B.L. & Bernardes, G.J.L. Contemporary approaches to site-selective protein modification. Nature Reviews Chemistry 3, 147–171 (2019).

14. Fantoni, N.Z., El-Sagheer, A.H. & Brown, T. A Hitchhiker’s Guide to Click-Chemistry with Nucleic Acids. Chemical Reviews 121, 7122–7154 (2021).

15. Lang, K. & Chin, J.W. Bioorthogonal Reactions for Labeling Proteins. ACS Chemical Biology 9, 16–20 (2014).

16. Keppler, A. et al. Labeling of fusion proteins of O6-alkylguanine-DNA alkyltransferase with small molecules in vivo and in vitro. Methods 32, 437–444 (2004).

17. Keppler, A., Pick, H., Arrivoli, C., Vogel, H. & Johnsson, K. Labeling of fusion proteins with synthetic fluorophores in live cells. Proceedings of the National Academy of Sciences 101, 9955–9959 (2004).

18. Kühn, S. et al. SNAP-tag2 for faster and brighter protein labeling. Nature Chemical Biology (2025).

19. Wolfram-Schauerte, M. et al. A viral ADP-ribosyltransferase attaches RNA chains to host proteins. Nature 620, 1054–1062 (2023).

20. Byrgazov, K. et al. Structural basis for the interaction of protein S1 with the Escherichia coli ribosome. Nucleic Acids Research 43, 661–673 (2014).

21. Diedrich, G. et al. Ribosomal protein L2 is involved in the association of the ribosomal subunits, tRNA binding to A and P sites and peptidyl transfer. Embo j 19, 5241–5250 (2000).

22. Draper, D.E., Pratt, C.W. & von Hippel, P.H. Escherichia coli ribosomal protein S1 has two polynucleotide binding sites. Proc Natl Acad Sci U S A 74, 4786–4790 (1977).

23. Nakagawa, A. et al. The three-dimensional structure of the RNA-binding domain of ribosomal protein L2; a protein at the peptidyl transferase center of the ribosome. The EMBO Journal 18, 1459–1467 (1999).

24. Cahová, H., Winz, M.-L., Höfer, K., Nübel, G. & Jäschke, A. NAD captureSeq indicates NAD as a bacterial cap for a subset of regulatory RNAs. Nature 519, 374–377 (2015).

25. Chen, Y.G., Kowtoniuk, W.E., Agarwal, I., Shen, Y. & Liu, D.R. LC/MS analysis of cellular RNA reveals NAD-linked RNA. Nat Chem Biol 5, 879–881 (2009).

26. Dong, H. et al. NAD(+)-capped RNAs are widespread in rice (Oryza sativa) and spatiotemporally modulated during development. Sci China Life Sci 65, 2121–2124 (2022).

27. Frindert, J. et al. Identification, Biosynthesis, and Decapping of NAD-Capped RNAs in B. subtilis. Cell Rep 24, 1890-1901.e1898 (2018).

28. Gomes-Filho, J. et al. Identification of NAD-RNAs and ADPR-RNA decapping in the archaeal model organisms Sulfolobus acidocaldarius and Haloferax volcanii. (2022).

29. Jiao, X. et al. 5’ End Nicotinamide Adenine Dinucleotide Cap in Human Cells Promotes RNA Decay through DXO-Mediated deNADding. Cell 168, 1015-1027.e1010 (2017).

30. Morales-Filloy, H.G. et al. The 5’ NAD Cap of RNAIII Modulates Toxin Production in Staphylococcus aureus Isolates. J Bacteriol 202 (2020).

31. Olatz, R.-L. et al. NAD+ capping of RNA in Archaea and Mycobacteria. bioRxiv, 2021.2012.2014.472595 (2021).

32. Walters, R.W. et al. Identification of NAD+ capped mRNAs in Saccharomyces cerevisiae. Proc Natl Acad Sci U S A 114, 480–485 (2017).

33. Wang, J. et al. Quantifying the RNA cap epitranscriptome reveals novel caps in cellular and viral RNA. Nucleic Acids Res 47, e130 (2019).

34. Wang, Y. et al. NAD(+)-capped RNAs are widespread in the Arabidopsis transcriptome and can probably be translated. Proc Natl Acad Sci U S A 116, 12094–12102 (2019).

35. Zhang, H. et al. NAD tagSeq reveals that NAD+-capped RNAs are mostly produced from a large number of protein-coding genes in Arabidopsis. Proc Natl Acad Sci U S A 116, 12072–12077 (2019).

36. Zhang, Y. et al. Extensive 5′-surveillance guards against non-canonical NAD-caps of nuclear mRNAs in yeast. Nature Communications 11, 5508 (2020).

37. Giraud, P., Créchet, J.-B., Uzan, M., Bontems, F. & Sizun, C. Resonance assignment of the ribosome binding domain of E. coli ribosomal protein S1. Biomolecular NMR Assignments 9, 107–111 (2015).

38. Keeble, A.H. et al. Approaching infinite affinity through engineering of peptide–protein interaction. Proceedings of the National Academy of Sciences 116, 26523–26533 (2019).

39. Tiemann, B., Depping, R. & Rüger, W. Overexpression, purification, and partial characterization of ADP-ribosyltransferases modA and modB of bacteriophage T4. Gene Expr 8, 187–196 (1999).

40. Cohen, M.S. & Chang, P. Insights into the biogenesis, function, and regulation of ADP-ribosylation. Nat Chem Biol 14, 236–243 (2018).

41. Jiang, H., Kim, J.H., Frizzell, K.M., Kraus, W.L. & Lin, H. Clickable NAD analogues for labeling substrate proteins of poly(ADP-ribose) polymerases. J Am Chem Soc 132, 9363–9372 (2010).

42. Wang, Y., Rösner, D., Grzywa, M. & Marx, A. Chain-terminating and clickable NAD^+^ analogues for labeling the target proteins of ADP-ribosyltransferases. Angew Chem Int Ed Engl 53, 8159–8162 (2014).

43. Krska, D., Ravulapalli, R., Fieldhouse, R.J., Lugo, M.R. & Merrill, A.R. C3larvin toxin, an ADP-ribosyltransferase from Paenibacillus larvae. J Biol Chem 290, 1639–1653 (2015).

44. Zhang, J. Use of biotinylated NAD to label and purify ADP-ribosylated proteins. Methods Enzymol 280, 255–265 (1997).

45. Davis, R.E. et al. In Situ Staining for Poly(ADP-Ribose) Polymerase Activity Using an NAD Analogue. Journal of Histochemistry & Cytochemistry 46, 1279–1289 (1998).

46. Depaix, A. & Kowalska, J. NAD Analogs in Aid of Chemical Biology and Medicinal Chemistry. Molecules 24, 4187 (2019).

47. Abramson, J. et al. Accurate structure prediction of biomolecular interactions with AlphaFold 3. Nature 630, 493–500 (2024).

48. Tiemann, B. et al. ModA and ModB, two ADP-ribosyltransferases encoded by bacteriophage T4: catalytic properties and mutation analysis. J Bacteriol 186, 7262–7272 (2004).

49. Höfer, K., Abele, F., Schlotthauer, J. & Jäschke, A. Synthesis of 5′-NAD-Capped RNA. Bioconjugate Chemistry 27, 874–877 (2016).

50. Li, L., Fierer, J.O., Rapoport, T.A. & Howarth, M. Structural analysis and optimization of the covalent association between SpyCatcher and a peptide Tag. J Mol Biol 426, 309–317 (2014).

51. Zakeri, B. et al. Peptide tag forming a rapid covalent bond to a protein, through engineering a bacterial adhesin. Proceedings of the National Academy of Sciences 109, E690–E697 (2012).

52. Rack, J.G.M. et al. (ADP-ribosyl)hydrolases: Structural Basis for Differential Substrate Recognition and Inhibition. Cell Chemical Biology 25, 1533-1546.e1512 (2018).

53. Jones, C.I., Zabolotskaya, M.V. & Newbury, S.F. The 5′ → 3′ exoribonuclease XRN1/Pacman and its functions in cellular processes and development. WIREs RNA 3, 455–468 (2012).

54. Fan, Q. et al. A review of conjugation technologies for antibody drug conjugates. Antib Ther 8, 157–170 (2025).

55. Tsui, N.B., Ng, E.K. & Lo, Y.D. Stability of Endogenous and Added RNA in Blood Specimens, Serum, and Plasma. Clinical Chemistry 48, 1647–1653 (2002).

56. Wang, C. & Liu, H. Factors influencing degradation kinetics of mRNAs and half-lives of microRNAs, circRNAs, lncRNAs in blood in vitro using quantitative PCR. Scientific Reports 12, 7259 (2022).

57. Yang, L., Li, Y., Bhattacharya, A. & Zhang, Y. A plasma proteolysis pathway comprising blood coagulation proteases. Oncotarget 7, 40919–40938 (2016).

58. Ebrahimi, S.B. & Samanta, D. Engineering protein-based therapeutics through structural and chemical design. Nature Communications 14, 2411 (2023).

59. Elliott, S. et al. Enhancement of therapeutic protein in vivo activities through glycoengineering. Nat Biotechnol 21, 414–421 (2003).

60. Harris, J.M. & Chess, R.B. Effect of pegylation on pharmaceuticals. Nature Reviews Drug Discovery 2, 214–221 (2003).

61. Pyzik, M., Kozicky, L.K., Gandhi, A.K. & Blumberg, R.S. The therapeutic age of the neonatal Fc receptor. Nature Reviews Immunology 23, 415–432 (2023).

62. Andries, O. et al. N(1)-methylpseudouridine-incorporated mRNA outperforms pseudouridine-incorporated mRNA by providing enhanced protein expression and reduced immunogenicity in mammalian cell lines and mice. J Control Release 217, 337–344 (2015).

63. Karikó, K., Buckstein, M., Ni, H. & Weissman, D. Suppression of RNA Recognition by Toll-like Receptors: The Impact of Nucleoside Modification and the Evolutionary Origin of RNA. Immunity 23, 165–175 (2005).

64. Karikó, K. et al. Incorporation of pseudouridine into mRNA yields superior nonimmunogenic vector with increased translational capacity and biological stability. Mol Ther 16, 1833–1840 (2008).

65. Sun, X., Setrerrahmane, S., Li, C., Hu, J. & Xu, H. Nucleic acid drugs: recent progress and future perspectives. Signal Transduction and Targeted Therapy 9, 316 (2024).

66. Paunovska, K., Loughrey, D. & Dahlman, J.E. Drug delivery systems for RNA therapeutics. Nature Reviews Genetics 23, 265–280 (2022).

67. Ma, Y., Li, S., Lin, X. & Chen, Y. A perspective of lipid nanoparticles for RNA delivery. Exploration 4, 20230147 (2024).

68. Riccardi, F., Dal Bo, M., Macor, P. & Toffoli, G. A comprehensive overview on antibody-drug conjugates: from the conceptualization to cancer therapy. Front Pharmacol 14, 1274088 (2023).

69. Nübel, G., Sorgenfrei, F.A. & Jäschke, A. Boronate affinity electrophoresis for the purification and analysis of cofactor-modified RNAs. Methods 117, 14–20 (2017).

70. Keeble, A.H. et al. Evolving Accelerated Amidation by SpyTag/SpyCatcher to Analyze Membrane Dynamics. Angew Chem Int Ed Engl 56, 16521–16525 (2017).

