## Supplementary Information for "Engineering site-specific nucleic acid-protein conjugates by utilizing a natural RNAylation reaction"



Supplementary Figure 1: pLDDT score of each residue on the AlphaFold3[47] prediction model created in PyMOL.
Supplementary Figure 2: Preparation of 5′ NXD-RNA 10mer Cy5.
Supplementary Figure 3: RNAylation of rS1 in the presence of 5′ NXD-RNA 10mers.
Supplementary Figure 4: Preparation of 5′ NXD-DNA 40mer Cy5.
Supplementary Figure 5: DNAylation of rS1 in the presence of 5′ NXD-DNA 40mers.
Supplementary Figure 6: RNAylation of rS1 DII in the presence of 5′ NXD-RNA 10mers.
Supplementary Figure 7: RNAylation of GFP enabled by the fusion of rS1 DII.
Supplementary Figure 8: Detected RNAylated residues on SpyTag-DII by LC-MS/MS analysis.
Supplementary Figure 9: *In vitro* ARH1 kinetics of NXD-RNAylated rS1.
Supplementary Figure 10: *In vitro* ARH1 kinetics of NXD-DNAylated rS1.
Supplementary Figure 11: *In vitro* XRN1 kinetics of NXD-RNAylated rS1.
Supplementary Figure 12: Human cell lysate stability kinetics of NXD-RNAylated GFP-rL2.
Supplementary Figure 13: Mouse blood plasma stability kinetics of NXD-RNAylated GFP-rL2.

Supplementary Figure 14: Purification of DNAylated GFP-rL2.
Supplementary Figure 15: Maximum intensity projection images of transfected HEK293T cells.
Supplementary Table 1: The sequences of 5′-monophosphorylated RNA and DNA oligos purchased from IDT.
Supplementary Table 2: The primer sequences to generate plasmids used in this study to produce target proteins.

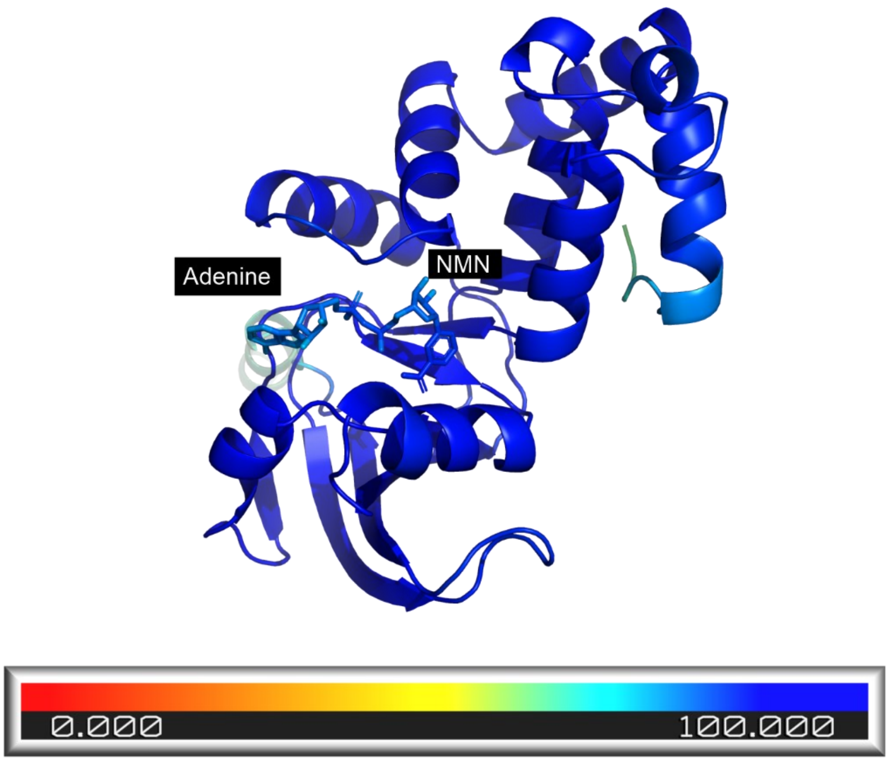


**Supplementary Figure 1: pLDDT score of each residue on the AlphaFold3[47] prediction model created in PyMOL.** The structural prediction of ModB is performed via AlphaFold3 in default settings with a single copy of ModB protein and a single copy of the NAD substrate. The AlphaFold3 predicted structure exhibits a high overall confidence with a predicted Template Modeling (pTM) value of 0.89, indicating accurate domain packing and overall topology. The pTM score ranges from 0 to 1, where values ≥ 0.7 are considered high confidence in the global structural arrangement. This figure depicts the color coding of per-residue confidence, which is evaluated by using predicted Local Distance Difference Test (pLDDT) scores, revealing consistently high reliability across the entire model. The structure is uniformly colored blue, corresponding to pLDDT values ≥90, which depicts very high confidence. The pLDDT color scheming for the structural visualization is as follows: blue (≥90, very high confidence), cyan (70–89, confident), yellow (50–69, low confidence), and orange to red (<50, very low confidence).

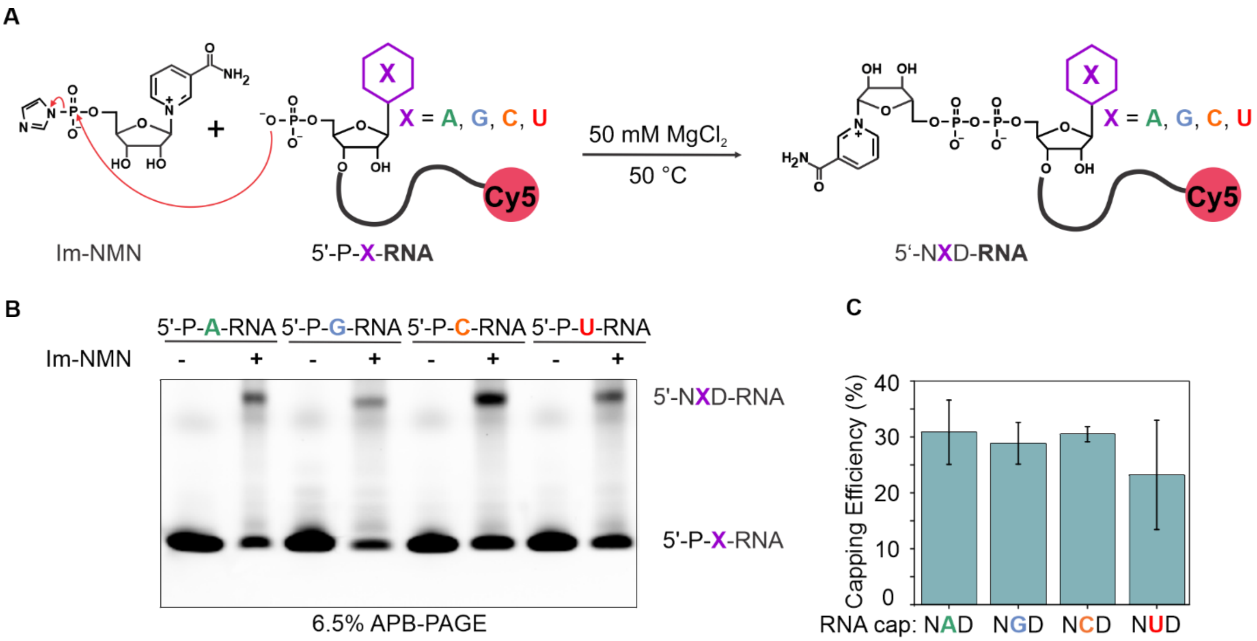


**Supplementary Figure 2: Preparation of 5′ NXD-RNA 10mer Cy5.** A) 5′-P-X-RNA 10mers were incubated in the presence of 1000-fold excess nicotinamide mononucleotide (NMN)-phosphorimidazolide (Im-NMN) to generate 5′ NXD-RNA by the coupling reaction of Im-NMN to the 5′-monophosphate group according to the published protocol[49]. B) Analysis of the 5′ NXD-capping was performed by analytical acryloylaminophenyl boronic acid (APB) gel electrophoresis[69]. 5′-P-X-RNA was run next to the 5′ NXD-RNA. C) The graph showing the comparison of capping efficiencies of 5′-NXD-RNA capping. Error bars represent the mean ± s.d. of n = 3 independent experiments.

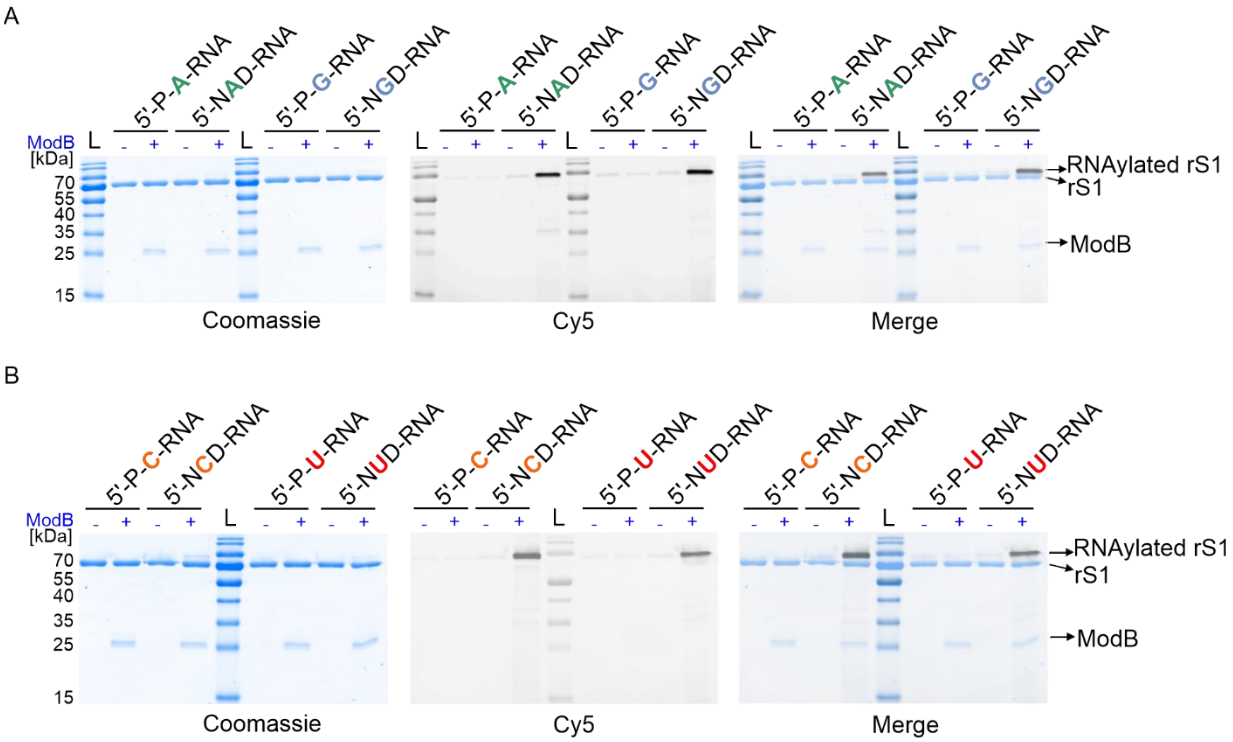


**Supplementary Figure 3: RNAylation of rS1 in the presence of 5′ NXD-RNA 10mers.** RNAylation reactions were analyzed by 12% SDS-PAGE. RNAylated rS1 was detected as a shifted band in the presence of NXD-RNA and ModB in the Cy5 channel for all tested NXD-RNA substrates. RNAylation is performed in the presence of 3′ Cy5-labelled NXD-RNA 10mers. (n = 3 independent experiments) A) The gel shows the analysis of RNAylation reactions in the presence of P-A-, NAD-, P-G-, and NGD-capped RNAs. B) The gel shows the analysis of RNAylation reactions in the presence of P-C-, NCD-, P-U-, and NUD-capped RNAs.

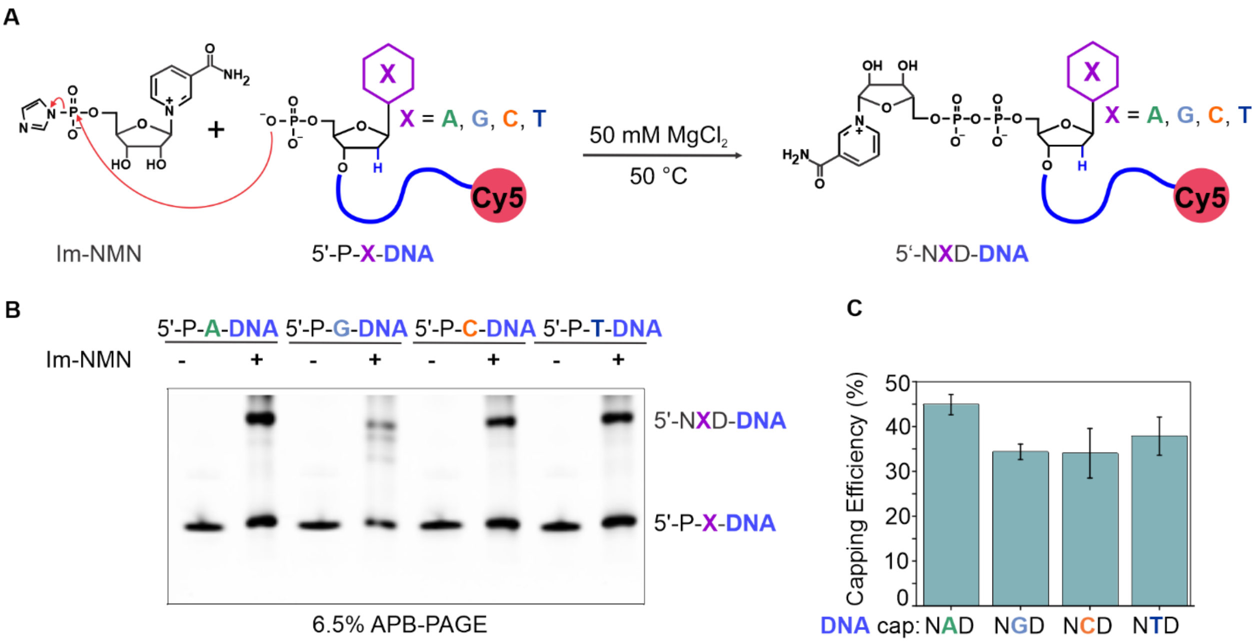


**Supplementary Figure 4: Preparation of 5′ NXD-DNA 40mer Cy5.** A) 5′-P-X-DNA 40mers were incubated in the presence of 1000-fold excess Im-NMN to generate 5′ NXD-DNA by the coupling reaction of Im-NMN to the 5′-monophosphate group according to the published protocol[49]. B) Analysis of the 5′ NXD-capping was performed by analytical APB gel electrophoresis. 5′-P-X-DNA was run next to the 5′ NXD-DNA. C) The graph showing the comparison of capping efficiencies of 5′ NXD-DNA capping. Error bars represent the mean ± s.d. of n = 3 independent experiments.

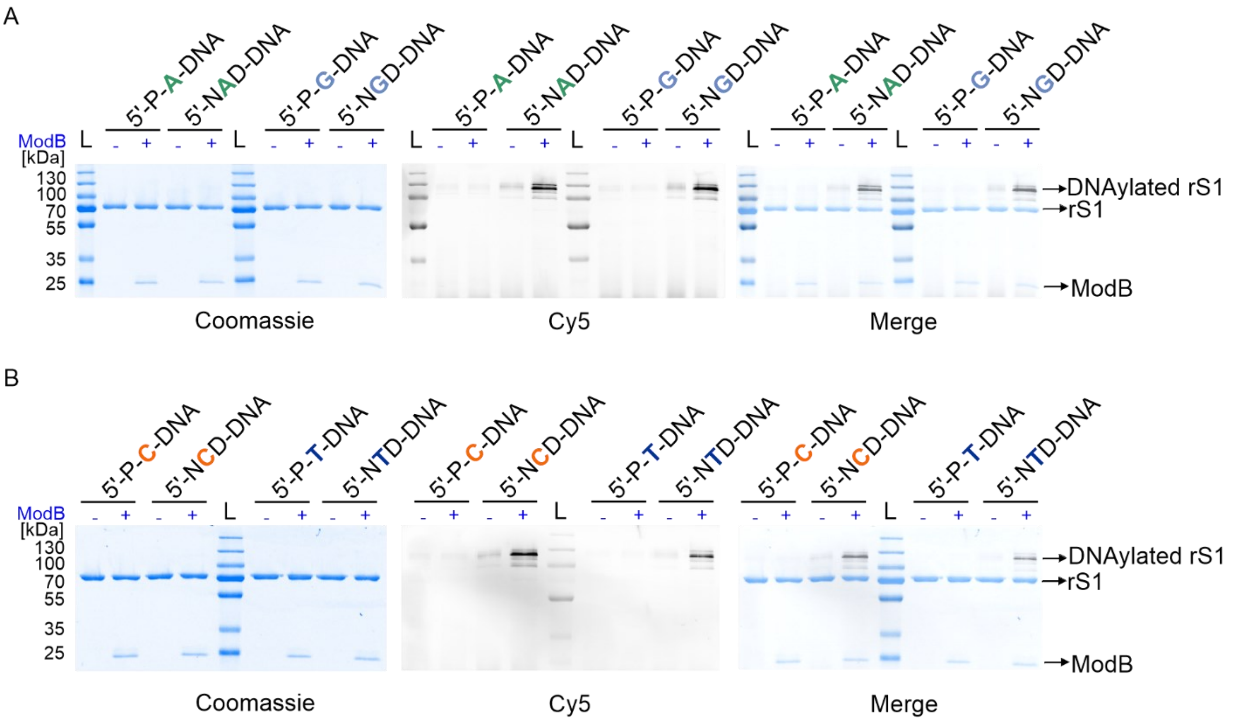


**Supplementary Figure 5: DNAylation of rS1 in the presence of 5′ NXD-DNA 40mers.** DNAylation reactions were analyzed by 12% SDS-PAGE. DNAylated rS1 was detected as a shifted band in the presence of NXD-DNA and ModB in the Cy5 channel for all tested NXD-DNA substrates. DNAylation reactions were performed in the presence of 3′ Cy5-labelled NXD-DNA 40mers. (n = 3 independent experiments) A) The gel shows the analysis of DNAylation reactions in the presence of P-A-, NAD-, P-G-, and NGD-capped DNAs. B) The gel shows the analysis of DNAylation reactions in the presence of P-C-, NCD-, P-T-, and NTD-capped DNAs.

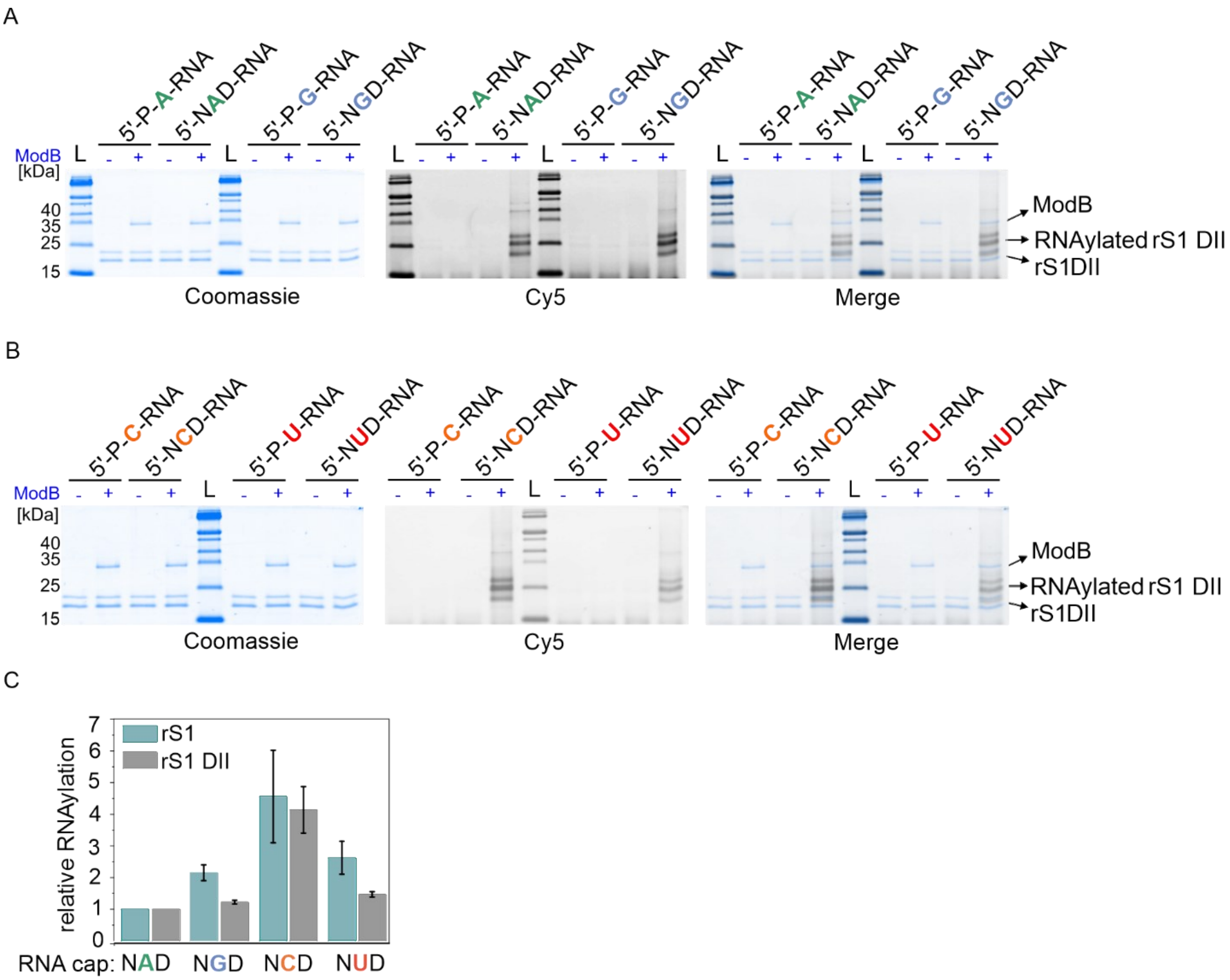


**Supplementary Figure 6: RNAylation of rS1 DII in the presence of 5′ NXD-RNA 10mers.** RNAylation reactions were analyzed by 15% Tricine-PAGE. RNAylated rS1 DII was detected as a shifted band in the presence of NXD-RNA and ModB in the Cy5 channel for all tested NXD-RNA substrates. A) The gel shows the analysis of RNAylation reactions in the presence of P-A-, NAD-, P-G-, and NGD-capped RNAs. B) The gel shows the analysis of RNAylation reactions in the presence of P-C-, NCD-, P-U-, and NUD-capped RNAs. C) The comparison of relative RNAylation efficiencies of rS1 and rS1 DII. Error bars represent the mean ± s.d. of n = 3 independent experiments.

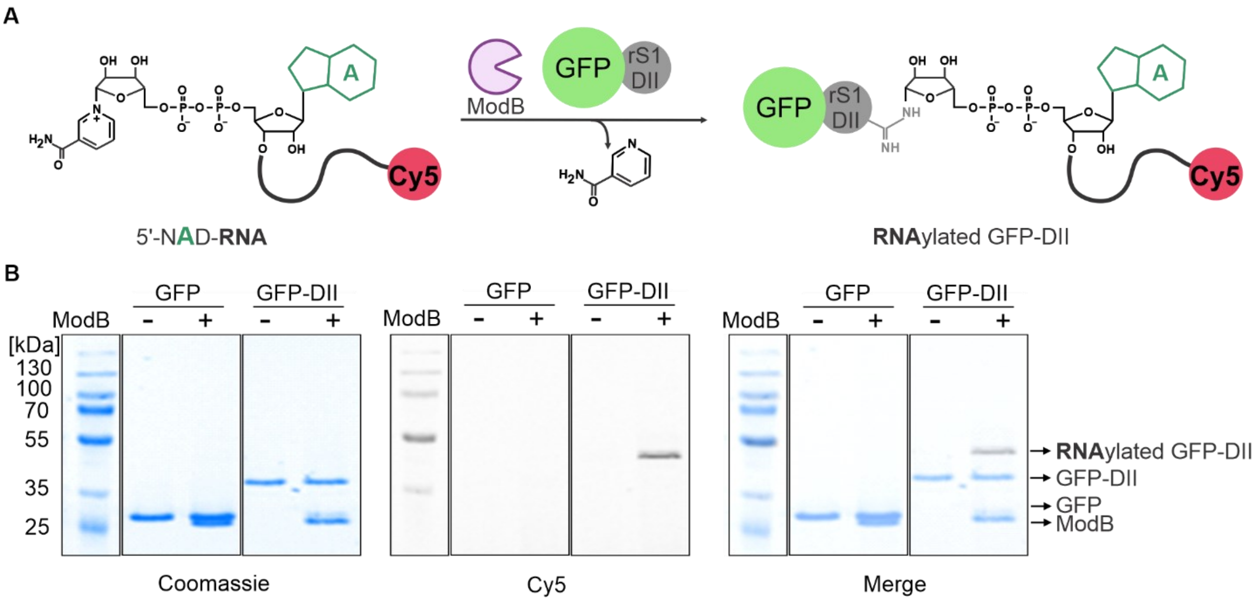


**Supplementary Figure 7: RNAylation of GFP enabled by the fusion of rS1 DII.** A) Proposed RNAylation reaction of GFP protein fused with rS1 DII (GFP-DII) in the presence of NAD-RNA. B) RNAylation reaction in the presence of 3′ Cy5-labeled NAD-RNA 10mer, GFP, and GFP-DII proteins in the presence and absence of ModB has been visualized by 12% SDS-PAGE. (n = 3 independent experiments)

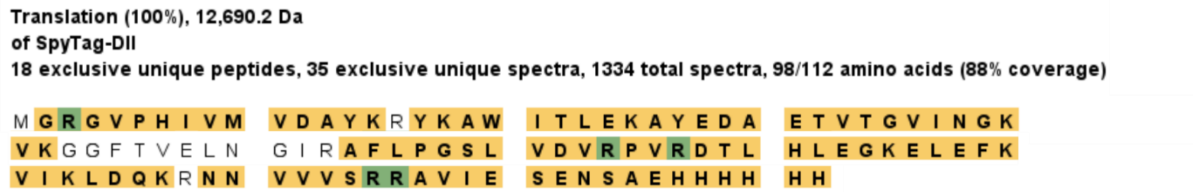


**Supplementary Figure 8: Detected RNAylated residues on SpyTag-DII by LC-MS/MS analysis.** The sample coverage is 88% where detected peptide fragments are highlighted in yellow. Moreover, arginine residues, which are RNAylated by ModB, are highlighted in green color. (n = 3 independent experiments)

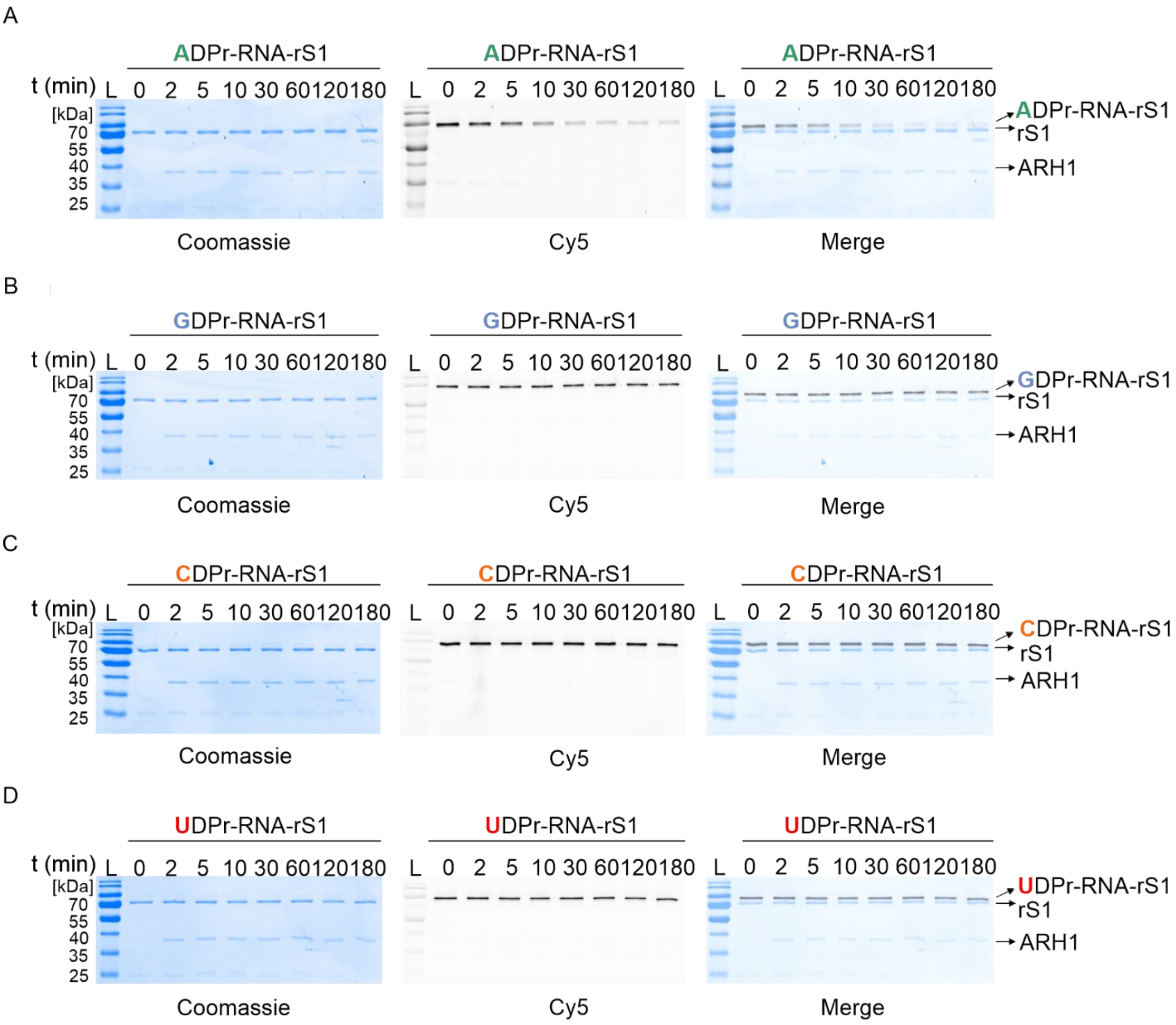


**Supplementary Figure 9: *In vitro* ARH1 kinetics of NXD-RNAylated rS1.** For RNAylation reactions, 3′ Cy5-labelled NXD-RNA 10mers have been utilized. RNAylated ADPr-RNA-rS1 (A), GDPr-RNA-rS1 (B), CDPr-RNA-rS1 (C), or UDPr-RNA-rS1 (D) proteins were subjected to ARH1 for 0, 2, 5, 10, 30, 60, 120, and 180 min. Reactions were analyzed by 12% SDS-PAGE. (n = 3 independent experiments)

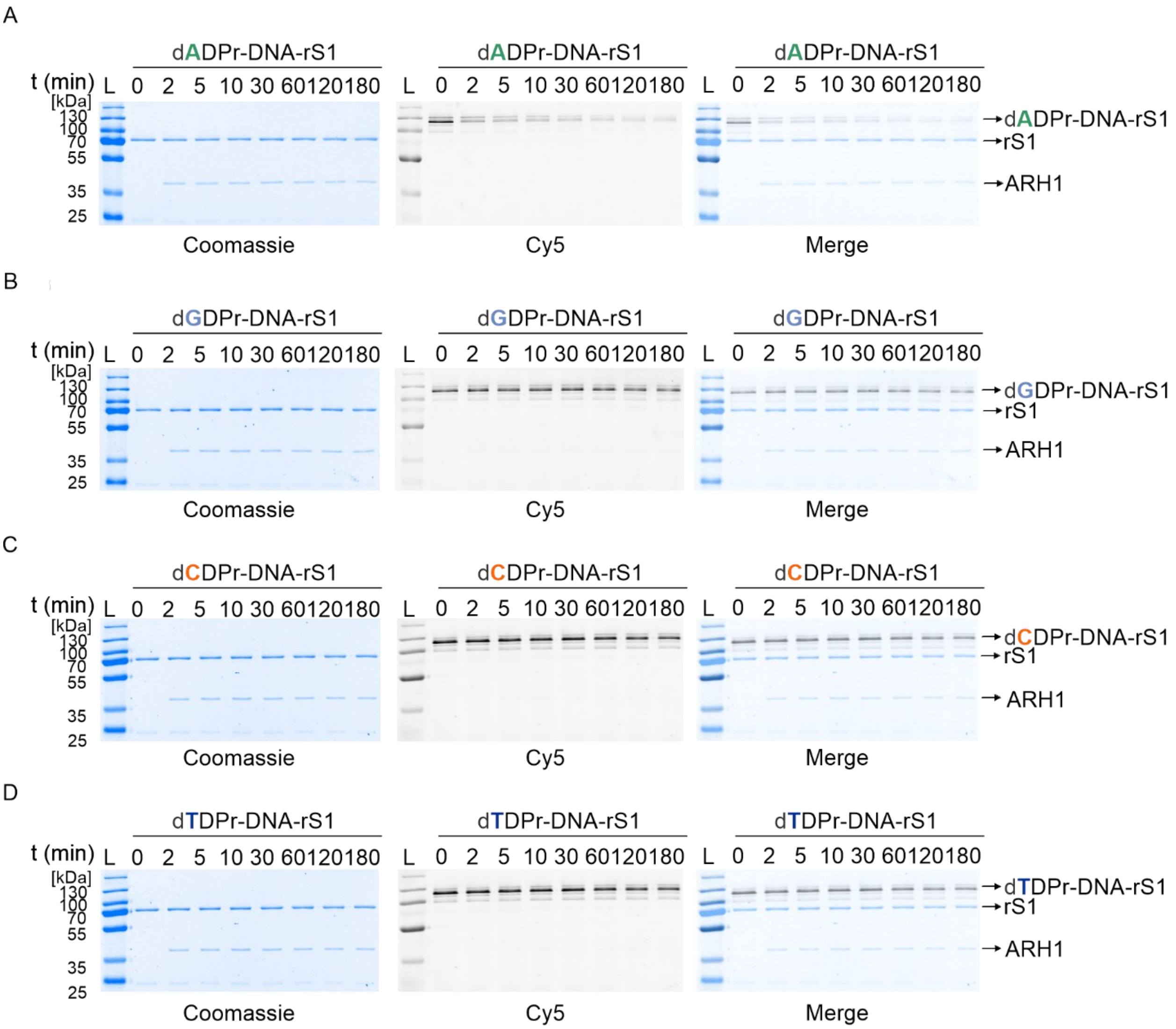


**Supplementary Figure 10: *In vitro* ARH1 kinetics of NXD-DNAylated rS1.** For DNAylation reactions, 3′ Cy5-labelled NXD-DNA 40mers were used. DNAylated ADPr-DNA-rS1 (A), GDPr-DNA-rS1 (B), CDPr-DNA-rS1 (C), or TDPr-DNA-rS1 (D) proteins were subjected to ARH1 for 0, 2, 5, 10, 30, 60, 120, and 180 min. Reactions were analyzed by 12% SDS-PAGE. (n = 3 independent experiments)

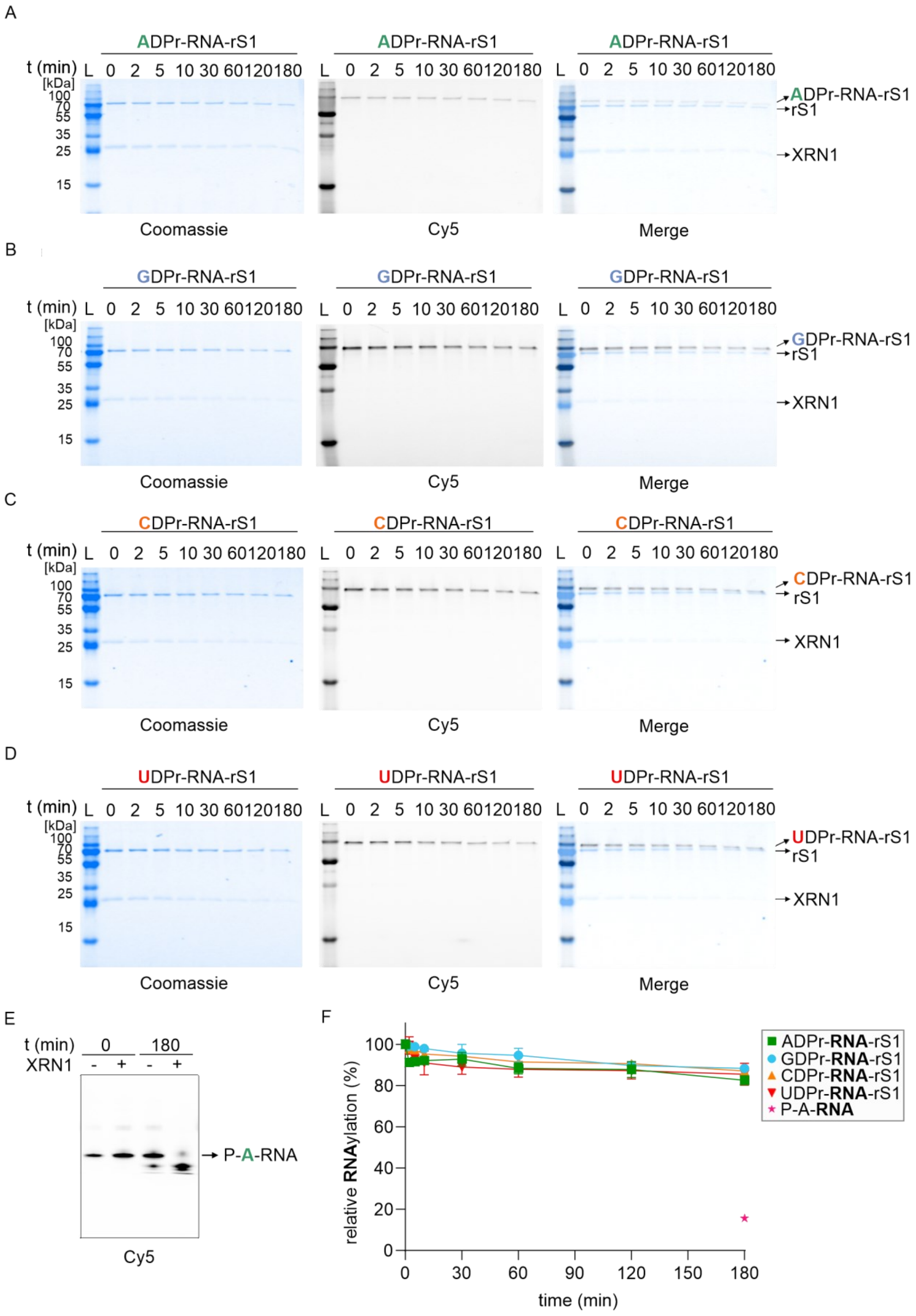
A
ADPr-RNA-rS1
t (min) L 0 2 5 10 30 60 120 180
[kDa]
100
70
55
35
25
15
ADPr-RNA-rS1
rS1
XRN1
Coomassie
Cy5
Merge
B
GDPr-RNA-rS1
GDPr-RNA-rS1
rS1
XRN1
C
CDPr-RNA-rS1
CDPr-RNA-rS1
rS1
XRN1
D
UDPr-RNA-rS1
UDPr-RNA-rS1
rS1
XRN1
E
t (min) 0 180
XRN1 - + - +
P-A-RNA
Cy5
F
relative RNAylation (%)
0
20
40
60
80
100
time (min)
0 30 60 90 120 150 180
ADPr-RNA-rS1
GDPr-RNA-rS1
CDPr-RNA-rS1
UDPr-RNA-rS1
P-A-RNA

**Supplementary Figure 11: *In vitro* XRN1 kinetics of NXD-RNAylated rS1.** For RNAylation reactions, 3′ Cy5-labelled NXD-RNA 10mers have been utilized. RNAylated ADPr-RNA-rS1 (A), GDPr-RNA-rS1 (B), CDPr-RNA-rS1 (C), or UDPr-RNA-rS1 (D) proteins were subjected to XRN1 for 0, 2, 5, 10, 30, 60, 120, and 180 min. Reactions were analyzed by 12% SDS-PAGE. E) P-A-RNA stability against XRN1 is analyzed by 20% PAGE. F) XRN1 kinetics graph is demonstrated. Error bars represent the mean ± s.d. of n = 3 independent experiments.

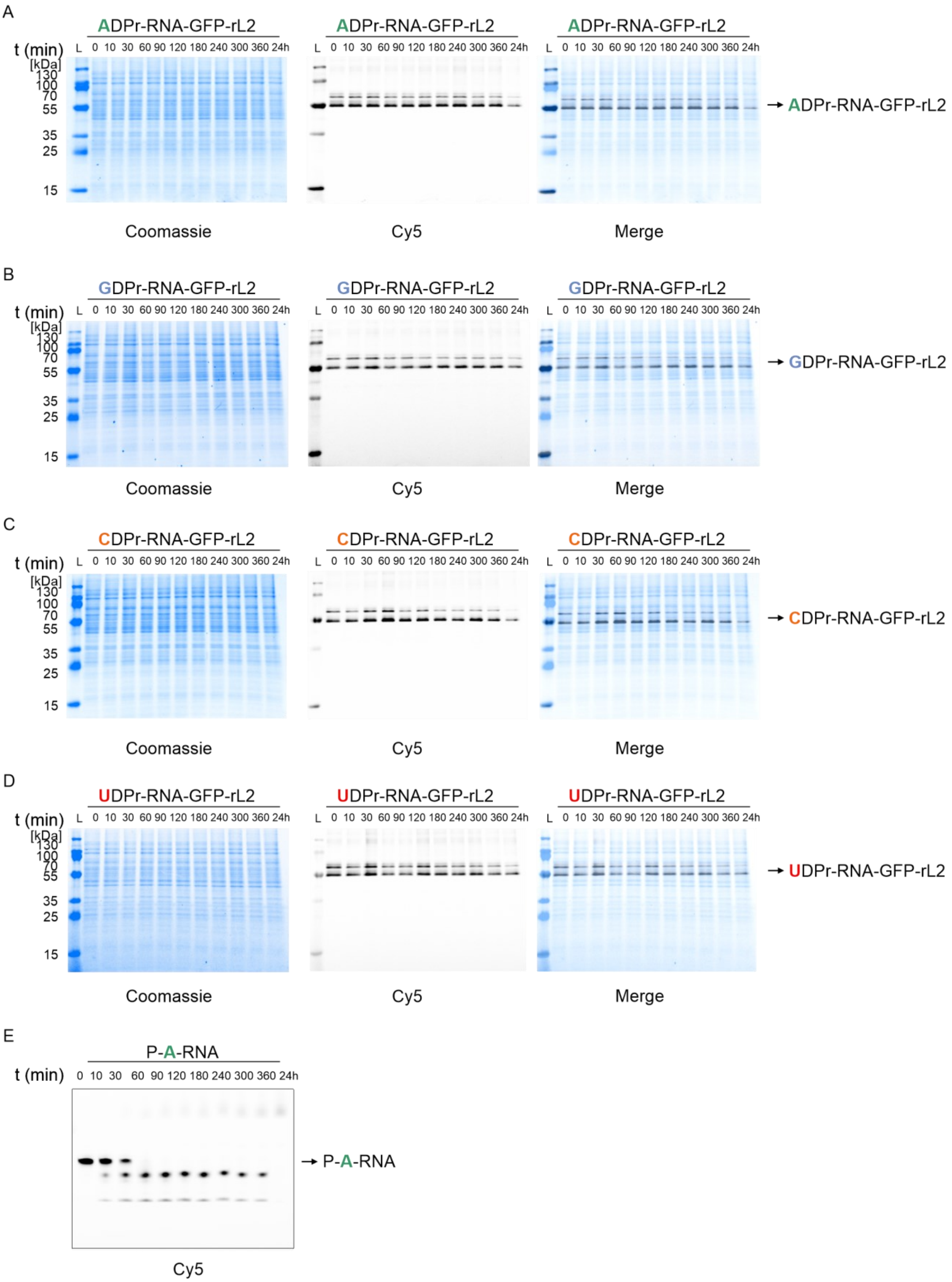
A
ADPr-RNA-GFP-rL2
t (min)
L 0 10 30 60 90 120 180 240 300 360 24h
[kDa]
130
100
70
55
35
25
15
→ ADPr-RNA-GFP-rL2
Coomassie
Cy5
Merge
B
GDPr-RNA-GFP-rL2
t (min)
L 0 10 30 60 90 120 180 240 300 360 24h
[kDa]
130
100
70
55
35
25
15
→ GDPr-RNA-GFP-rL2
Coomassie
Cy5
Merge
C
CDPr-RNA-GFP-rL2
t (min)
L 0 10 30 60 90 120 180 240 300 360 24h
[kDa]
130
100
70
55
35
25
15
→ CDPr-RNA-GFP-rL2
Coomassie
Cy5
Merge
D
UDPr-RNA-GFP-rL2
t (min)
L 0 10 30 60 90 120 180 240 300 360 24h
[kDa]
130
100
70
55
35
25
15
→ UDPr-RNA-GFP-rL2
Coomassie
Cy5
Merge
E
P-A-RNA
t (min)
0 10 30 60 90 120 180 240 300 360 24h
→ P-A-RNA
Cy5

**Supplementary Figure 12: Human cell lysate stability kinetics of NXD-RNAylated GFP-rL2.** For RNAylation reactions, 3′ Cy5-labelled NXD-RNA 10mers have been utilized. RNAylated ADPr-RNA-GFP-rL2 (A), GDPr-RNA-GFP-rL2 (B), CDPr-RNA-GFP-rL2 (C), or UDPr-RNA-GFP-rL2 (D) proteins were incubated in HEK293T cell lysates for 0, 10, 30, 60, 90, 120,180, 240, 300, 360 minutes and 24 hours. Reactions were analyzed by 12% SDS-PAGE. E) P-A-RNA kinetics is performed as a positive control to show nuclease activity in the human cell lysate. Reactions were analyzed by 20% PAGE. (n = 3 independent experiments)

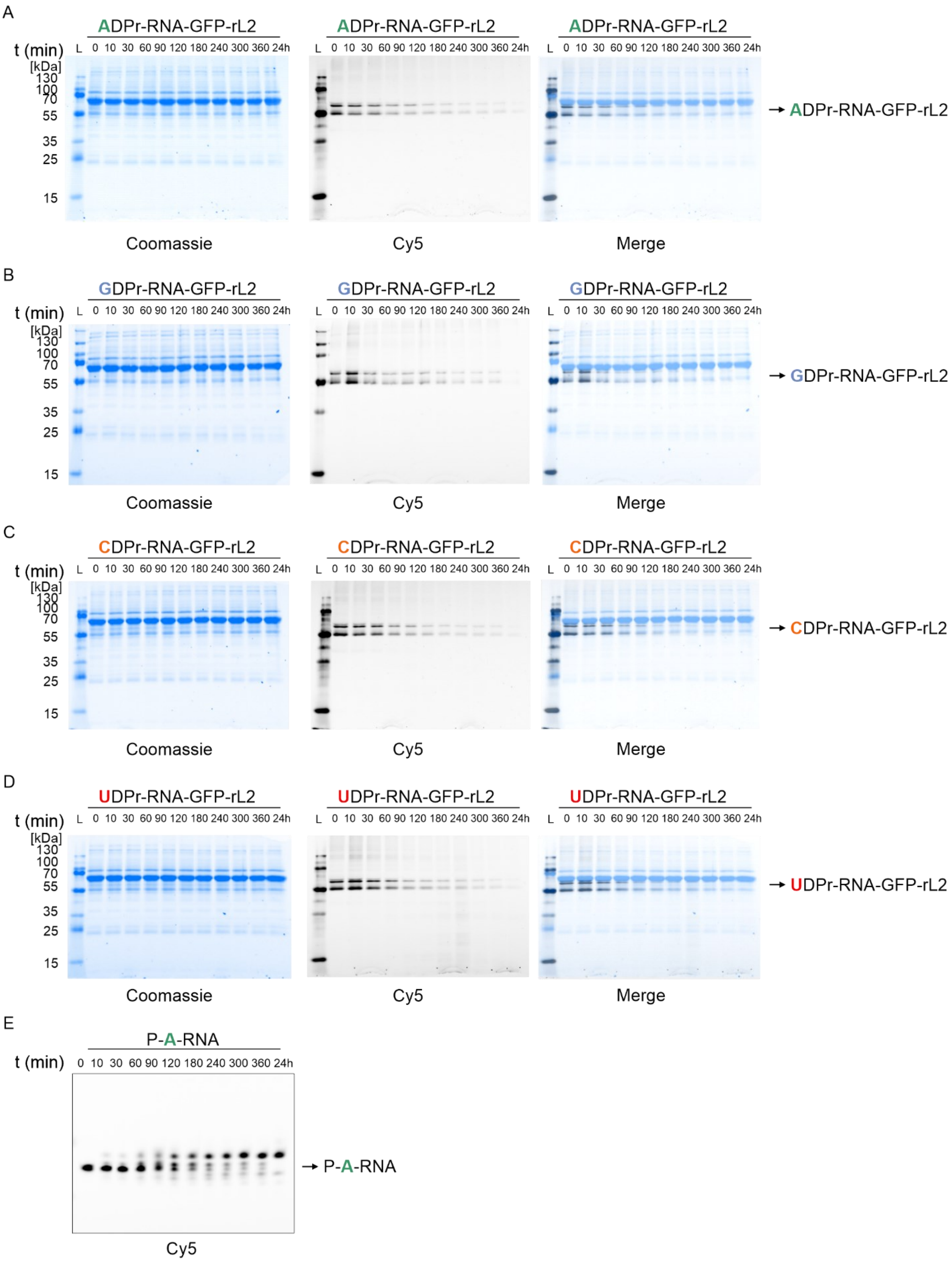
A
ADPr-RNA-GFP-rL2
t (min)
L 0 10 30 60 90 120 180 240 300 360 24h
[kDa]
130
100
70
55
35
25
15
Coomassie
Cy5
Merge
→ ADPr-RNA-GFP-rL2
B
GDPr-RNA-GFP-rL2
→ GDPr-RNA-GFP-rL2
C
CDPr-RNA-GFP-rL2
→ CDPr-RNA-GFP-rL2
D
UDPr-RNA-GFP-rL2
→ UDPr-RNA-GFP-rL2
E
P-A-RNA
t (min)
0 10 30 60 90 120 180 240 300 360 24h
→ P-A-RNA
Cy5

**Supplementary Figure 13: Mouse blood plasma stability kinetics of NXD-RNAylated GFP-rL2.** For RNAylation reactions, 3' Cy5-labelled NXD-RNA 10mers have been utilized. RNAylated ADPr-RNA-GFP-rL2 (A), GDPr-RNA-GFP-rL2 (B), CDPr-RNA-GFP-rL2 (C), or UDPr-RNA-GFP-rL2 (D) proteins were incubated in mouse blood plasma for 0, 10, 30, 60, 90, 120,180, 240, 300, 360 minutes and 24 hours. Reactions were analyzed by 12% SDS-PAGE. E) P-A-RNA kinetics is performed as a positive control to show nuclease activity in the blood plasma. Reactions were analyzed by 20% PAGE. (n = 3 independent experiments)

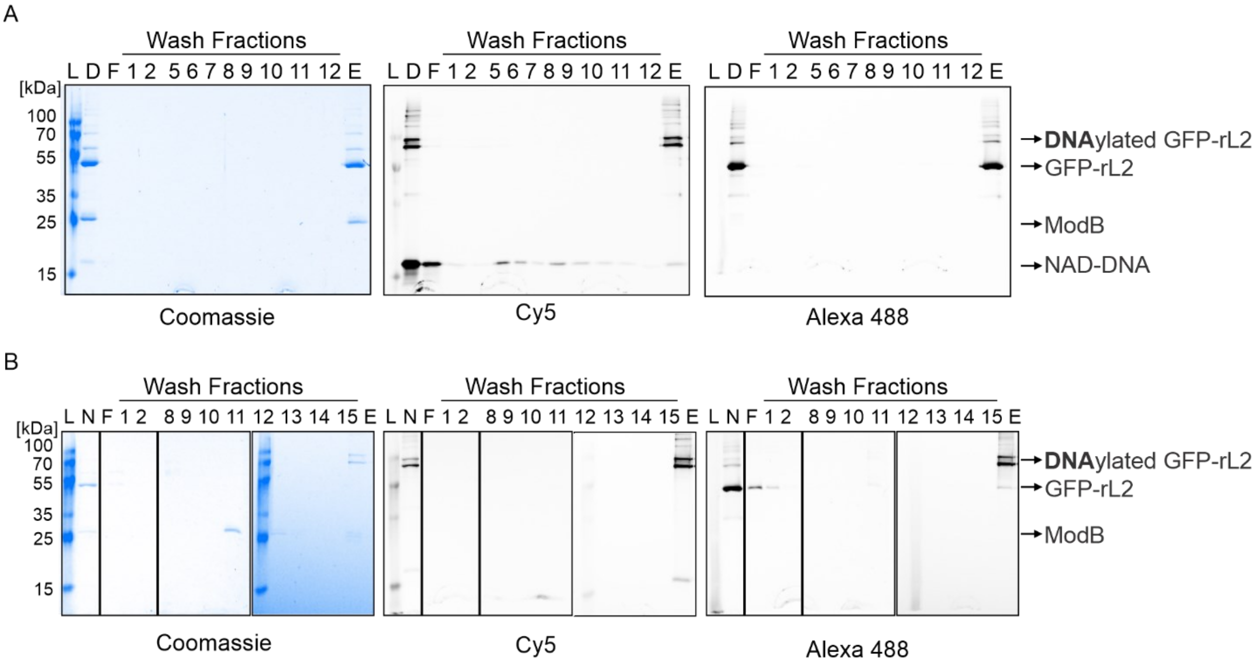


**Supplementary Figure 14: Purification of DNAylated GFP-rL2.** For DNAylation reactions, 3' Cy5-labelled NAD-DNA 60mer has been utilized. Purification products were analyzed by 12% SDS-PAGE. (L; Ladder, D: DNAylation reaction, F: Flow through, E: Elution, N: Denaturing IMAC purification product) (n = 3 independent experiments) A) Denaturing IMAC fractions were analyzed by 12% SDS-PAGE. B) Denaturing DEAE fractions were analyzed by 12% SDS-PAGE.

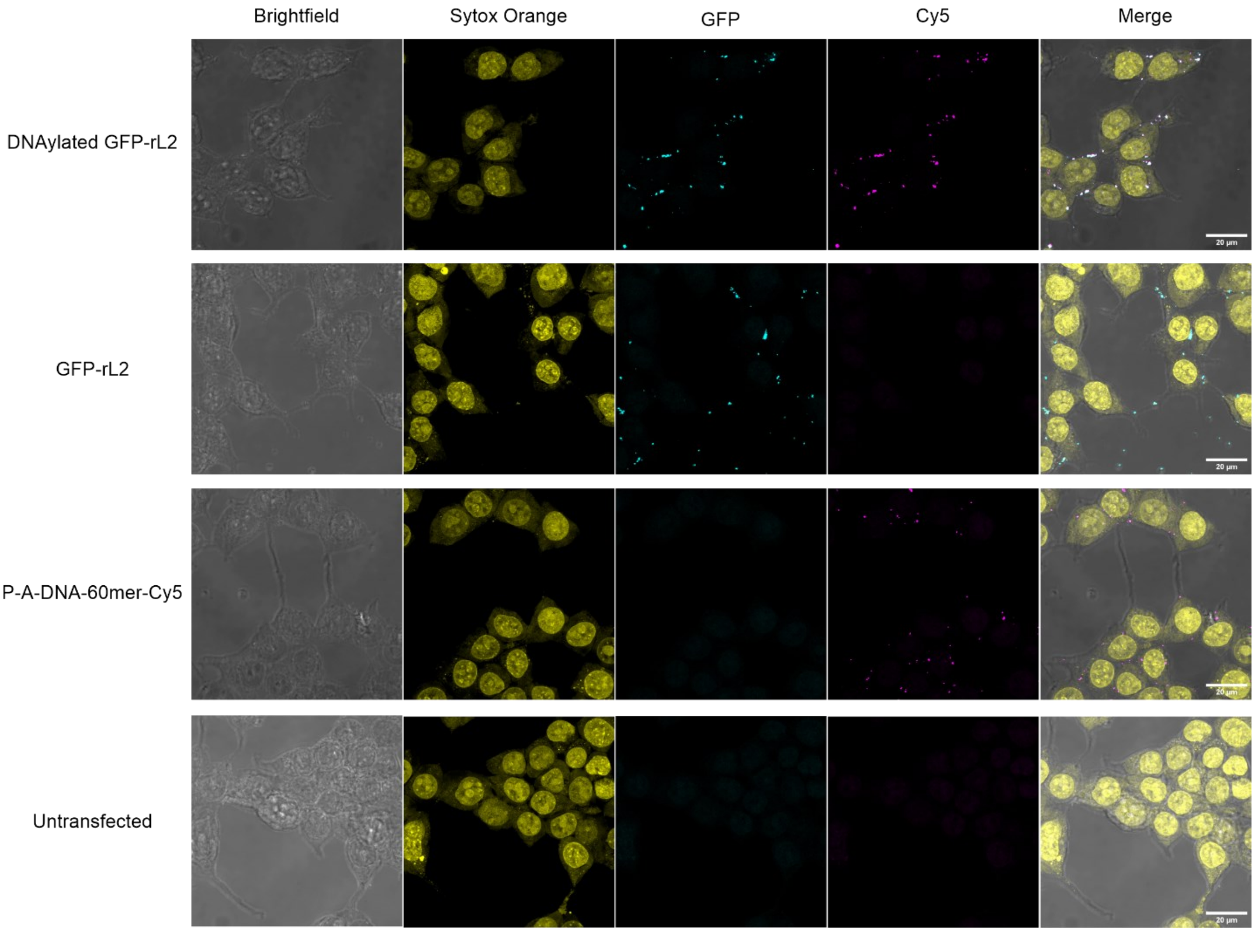


**Supplementary Figure 15: Maximum intensity projection images of transfected HEK293T cells.** In addition to purified DNAylated GFP-rL2 (DNAylation is performed with 3’Cy5 labelled NAD-DNA 60mer) delivered HEK293T cells, 5’-P-A-DNA 60mer Cy5, GFP-rL2, and untransfected controls are performed (n = 3 independent experiments). All samples were stained with Sytox Orange to visualize nuclei. For each sample, upon detecting the most focused z-plane, +/- 10 z-planes have been subjected to maximum intensity projection analysis to generate the images. The single channels Brightfield, Sytox Orange staining, GFP, Cy5, and the resulting merged images are shown with a representative image panel for each sample.

**Supplementary Table 1: The sequences of 5′-monophosphorylated RNA and DNA oligos purchased from IDT.** /5Phos/ represents the 5′-monophosphorylation and /3Cy5Sp/ shows 3′ Cy5 labeling of oligos for visualization purposes. The sequences are given in the direction of 5′ to 3′.

| Oligo name | Sequence |
|---|---|
| 5′-P-A-RNA 10mer Cy5 | /5Phos/AGCAUUCGAC/3Cy5Sp/ |
| 5′-P-G-RNA 10mer Cy5 | /5Phos/GGCAUUCGAC/3Cy5Sp/ |
| 5′-P-C-RNA 10mer Cy5 | /5Phos/CGCAUUCGAC/3Cy5Sp/ |
| 5′-P-U-RNA 10mer Cy5 | /5Phos/UGCAUUCGAC/3Cy5Sp/ |
| 5′-P-A-DNA 40mer Cy5 | /5Phos/ACAGTATTTGGTATCTGCGCTCTGCTGAAGCCAGTTACCT /3Cy5Sp/ |
| 5′-P-G-DNA 40mer Cy5 | /5Phos/GCAGTATTTGGTATCTGCGCTCTGCTGAAGCCAGTTACCT /3Cy5Sp/ |
| 5′-P-C-DNA 40mer Cy5 | /5Phos/CCAGTATTTGGTATCTGCGCTCTGCTGAAGCCAGTTACCT /3Cy5Sp/ |
| 5′-P-T-DNA 40mer Cy5 | /5Phos/TCAGTATTTGGTATCTGCGCTCTGCTGAAGCCAGTTACCT /3Cy5Sp/ |
| 5′-P-A-DNA 60mer Cy5 | /5Phos/ATCTTGATACTACCTTTAGTTCGTTTAAACACGTTCTTGATA GTATCTTTTTATTAACCC/3Cy5Sp/ |

**Supplementary Table 2: The primer sequences to generate plasmids used in this study to produce target proteins.** The primers are purchased from IDT. Some primers are 5′-phosphorylated (shown as /5Phos/ in the sequences) to self-circularize PCR products.

| Primer name | Sequence |
|---|---|
| *pET28a*-SpyTag-DII-$His_6$-Forward primer | /5Phos/ATGGTGGACGCCTACAAACGCTATAAAGCCTGGATCACG CTGGAAAAAGCTTACG |
| *pET28a*-SpyTag-DII-$His_6$-Reverse primer | /5Phos/AACAATATGAGGAACGCCACGTCCCATGGTATATCTCCTT CTTAAAGTTAAAACAAAATTA 3′ |
| *pET28a*-SpyTag R3K-DII-$His_6$- Forward primer | /5Phos/ATGGTGGACGCCTACAAACGCTATAAAGCCTGGATCACG CTGGAAAAAGCTTACG (the same as *pET28a*-SpyTag-DII-$His_6$-Forward primer) |
| *pET28a*-SpyTag R3K-DII-$His_6$- Reverse primer | /5Phos/AACAATATGAGGAACGCCCTTTCCCATGGTATATCTCCTTC TTAAAGTTAAAACAAAATT 3′ |
| *pET28a*-GFP-rL2-$His_6$-Forward primer (insert) | GTTTAACTTTAAGAAGGAGATATACCATGCGTAAAGGCGAAGAGCT GTTCACTGGTG |
| *pET28a*-GFP-rL2-$His_6$-Reverse primer (insert) | CGGTTTACATTTAACAACTGCGCCCATTCCAGCACCGGATCCTTTGTA CAGTTCATCCATACCATGCG |
| *pET28a*-GFP-rL2-$His_6$-Forward primer (backbone) | ATGGGCGCAGTTGTTAAATGTAAACCG |
| *pET28a*-GFP-rL2-$His_6$-Reverse primer (backbone) | GGTATATCTCCTTCTTAAAGTTAAAC |
| Inserted sfGFP sequence | ATGCGTAAAGGCGAAGAGCTGTTCACTGGTGTCGTCCCTATTCTGG TGGAACTGGATGGTGATGTCAACGGTCATAAGTTTTCCGTGCGTGG CGAGGGTGAAGGTGACGCAACTAATGGTAAACTGACGCTGAAGTT CATCTGTACTACTGGTAAACTGCCGGTACCTTGGCCGACTCTGGTA ACGACGCTGACTTATGGTGTTCAGTGCTTTGCTCGTTATCCGGACCA TATGAAGCAGCATGACTTCTTCAAGTCCGCCATGCCGGAAGGCTATG TGCAGGAACGCACGATTTCCTTTAAGGATGACGGCACGTACAAAACG CGTGCGGAAGTGAAATTTGAAGGCGATACCCTGGTAAACCGCATTGA GCTGAAAGGCATTGACTTTAAAGAAGACGGCAATATCCTGGGCCATA AGCTGGAATACAATTTTAACAGCCACAATGTTTACATCACCGCCGATA AACAAAAAAATGGCATTAAAGCGAATTTTAAAATTCGCCACAACGTG GAGGATGGCAGCGTGCAGCTGGCTGATCACTACCAGCAAAACACTC CAATCGGTGATGGTCCTGTTCTGCTGCCAGACAATCACTATCTGAGC ACGCAAAGCGTTCTGTCTAAAGATCCGAACGAGAAACGCGATCATA TGGTTCTGCTGGAGTTCGTAACCGCAGCGGGCATCACGCATGGTATG GATGAACTGTACAAAGGATCCGGTGCTGGA |
